# Defining the T cell transcriptional landscape in pediatric liver transplant rejection at single cell resolution

**DOI:** 10.1101/2024.02.26.582173

**Authors:** Anna L. Peters, Erica A.K. DePasquale, Gousia Begum, Krishna M. Roskin, E. Steve Woodle, David A. Hildeman

## Abstract

Acute cellular rejection (ACR) affects >80% of pediatric liver transplant recipients within 5 years, and late ACR is associated with graft failure. Traditional anti-rejection therapy for late ACR is ineffective and has remained unchanged for six decades. Although CD8+ T cells promote late ACR, little has been done to define their specificity and gene expression. Here, we used single-cell sequencing and immune repertoire profiling (10X Genomics) on 30 cryopreserved 16G liver biopsies from 14 patients (5 pre-transplant or with no ACR, 9 with ACR). We identified expanded intragraft CD8+ T cell clonotypes (CD8_EXP_) and their gene expression profiles in response to anti-rejection treatment. Notably, we found that expanded CD8^+^ clonotypes (CD8_EXP_) bore markers of effector and CD56^hi^CD161^-^ ‘NK-like’ T cells, retaining their clonotype identity and phenotype in subsequent biopsies from the same patients despite histologic ACR resolution. CD8_EXP_ clonotypes localized to portal infiltrates during active ACR, and persisted in the lobule after histologic ACR resolution. CellPhoneDB analysis revealed differential crosstalk between KC and CD8_EXP_ during late ACR, with activation of the LTB-LTBR pathway and downregulation of TGFß signaling. Therefore, persistently-detected intragraft CD8_EXP_ clones remain active despite ACR treatment and may contribute to long-term allograft fibrosis and failure of operational tolerance.

**Graphical Abstract:** 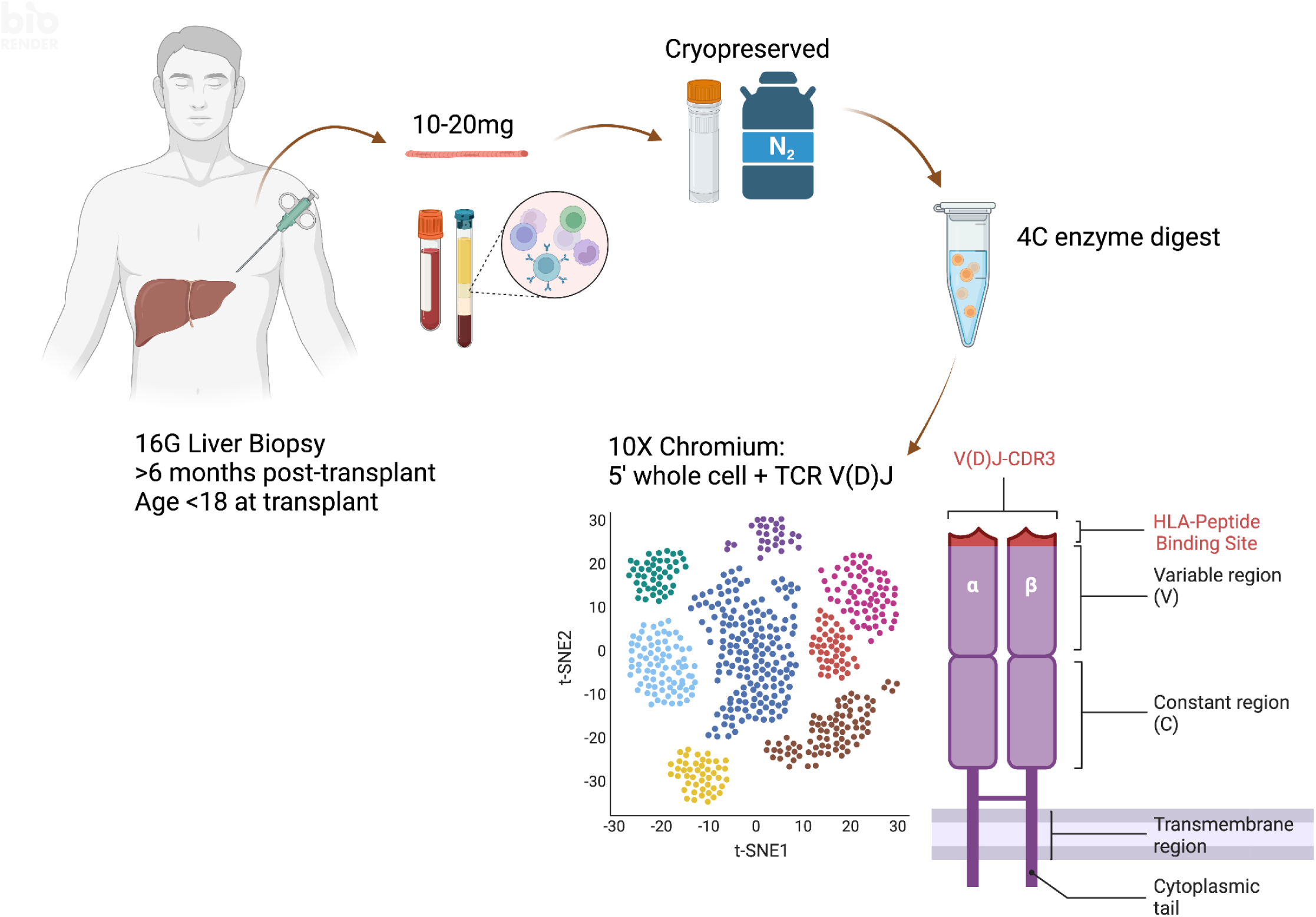

## Introduction

Liver transplantation remains the only treatment for end-stage liver disease (1, 2). Despite widespread adoption of tacrolimus as maintenance immunosuppression in pediatric liver transplantation (3), >50% of pediatric liver transplant recipients undergo acute cellular rejection (ACR) within the first post-transplant year (3–11). Of these ACR events, those occurring late (>6 months post-transplant) are more likely to lead to development of donor specific antibodies, chronic rejection, and graft loss in pediatric patients (6, 7, 12–17). Further, management of pediatric liver transplant recipients is challenging (4, 5). By nature of their young age, vulnerable growth period, and need for lifelong immunosuppression, pediatric patients are at increased risk for post-transplant lymphoproliferative disorder (PTLD), chronic kidney disease, and growth failure as compared to their adult counterparts (4, 5).

Studies of immunosuppression withdrawal in pediatric patients show that patients with occult allograft inflammation despite normal liver tests have a liver bulk gene expression profile similar to those undergoing ACR (18, 19), indicating that subclinical rejection can occur despite the presence of normal liver biochemistries. Further, increased CD8^+^ infiltrates and increased APC:CD8^+^ T cell immune synapses within the hepatic lobule (rather than portal areas) are predictive of operational tolerance. However, because these studies were done using bulk RNAseq, these genes are unable to be attributed to a specific cell type or cell-cell interaction (18–21). Although ACR is thought to be primarily mediated by CD8^+^ T cells (22–24), studies investigating the role of T cells in pediatric liver transplant rejection have been primarily limited to analysis of peripheral blood mononuclear cells (PBMC) (25, 26). Yet, the liver is an immune organ, housing an array of specialized innate immune cell populations (NK, NKT, Mucosal Associated Invariant T (MAIT)) in addition to cognate immune populations (CD4^+^ or CD8^+^ T cells) which are transplanted with the allograft into the recipient (27–31). To date, little has been done to investigate the signaling properties of intragraft T cells in liver transplant rejection beyond bulk RNA sequencing (18, 32, 33), tissue microarray (34), imaging mass cytometry (35), and TCR Vβ gene sequencing (36), which yield little insight into rejection pathogenesis beyond known pathways of antigen presentation to and interferon-ɣ production by T cells. A more detailed understanding of the underlying biology of rejection is needed, specifically in pediatric patients, to ensure long-term graft survival and minimize unintended consequences of immunosuppression.

Single cell RNA sequencing analysis of adult kidney allograft rejection has recently revealed that a limited number of expanded CD8^+^ T cell clones (CD8_EXP_) are found in the tissue and urine during ACR and that these expanded clones persist in the tissue after rejection treatment, changing their gene expression profiles in response to differing immunosuppressive regimens (37). Similar findings were recapitulated in bronchioalveolar lavage fluid from rejecting lung transplant recipients (38). These paradigm-shifting studies demonstrate that CD8_EXP_ alloreactive T cells are able to persist in solid organ allograft target tissue and adjacent biofluids, acquire markers associated with tissue residence and memory formation, and are resistant to treatment with anti-rejection protocols currently in use (37, 38). Although many high-quality single cell RNA sequencing studies have shown various T cell and macrophage subsets in the human liver primarily using freshly processed liver tissue (39–48), no studies have been performed to date specifically on pediatric liver transplant rejection or with cryopreserved specimens. Moreover, the dynamics of these populations following rejection therapy have also not been explored. As such, the relative contribution of clonally expanded intragraft CD8^+^ T cells to pediatric liver transplant rejection has yet to be elucidated.

Our previous work implicated increased CD8^+^ T cell infiltrates in portal areas as a marker for delayed rejection treatment response (49). Here, we characterized the CD8^+^ T cell infiltrates and cellular interacting partners in an unbiased manner, utilizing single cell RNA sequencing combined with TCR V(D)J sequencing of cryopreserved liver biopsy tissue obtained from rejecting vs. non-rejecting pediatric liver transplant recipients. We made the striking observation that increased CD8_EXP_ clonotypic infiltrates present during rejection persisted even in patients with biopsy-proven histologically resolved rejection, sometimes beyond 6 months after histologic and biochemical resolution of a rejection episode. Supervised gene expression analysis of CD8_EXP_ shows that these clones remain active, with increased expression of *HLA-DRA*, *GZMB*, and *CXCR6*. Based on their unique TCR sequence, we analyzed CD8_EXP_ clones in serial liver biopsies from the same patients, and demonstrated their localization in both portal and lobular areas of the liver. Interestingly, we uncovered evidence of macrophage-T cell crosstalk that implicated differential TNFR signaling pathways as a potential mechanism for clonal retention of CD8_EXP_ in liver tissue in recurrent and resolved ACR.

## Results

### Single-cell analysis of liver biopsies

To investigate T cell clonal expansion and gene expression changes in response to anti-rejection treatment in patients with ACR, we performed paired 5’ single-cell RNA and V(D)J sequencing (10X Genomics) on cells isolated from 30 liver biopsies obtained from 14 patients: 5 normal or non-ACR allograft dysfunction and 9 undergoing ACR. Pediatric transplant recipients varied by age (8 months-14 years), sex, race, and transplant indication, while donors varied in age, sex, race (**Table 1**). In the 11 patients with a post-transplant liver biopsy, 7 were transplanted for biliary atresia, as this is the most common indication for pediatric liver transplantation. To decrease the potential for recurrent autoimmunity confounding our analysis, no patients in this report were transplanted for autoimmune hepatitis, which can recur in the allograft after transplantation and share features which are difficult to histologically distinguish from ACR (10). Multiple biopsies were collected from 6 of the 14 patients post transplantation, and in 4 of these patients multiple biopsies were collected during ACR (**Table 2**). All of the biopsies were obtained ‘for cause’ due to abnormal liver biochemistries rather than via surveillance protocol (i.e. at pre-selected times post-transplant regardless of liver biochemistries). The alanine aminotransferase level (ALT), gamma glutamyl transferase level (GGT), and presence or absence of donor specific antibodies (DSA) measured at the time of each biopsy are detailed in **Table 2**. The biopsies were scored for rejection by a single blinded pathologist according to the Banff score (15), with no significant differences in Banff sub-scores between each rejection biopsy (data not shown). The majority of patients were on a baseline immunosuppression regimen of tacrolimus (TAC) at the time of initial study biopsy (**Table 2**). No significant differences were observed in the TAC levels between Late ACR and the Resolved ACR or No ACR control groups (**Table 2**).

**Table 1.** Patient demographics.

| Patient | Histologic Diagnosis | Prior ACR (Y/N) | Donor age/sex (years) | Recipient age/sex (years) | Donor race/ethnicity | Recipient race/ethnicity | Transplant indication |
| --- | --- | --- | --- | --- | --- | --- | --- |
| 1 | Pre-TXP | - | 15yo/M | - | Unknown | - | - |
| 2 | Pre-TXP | - | 46yo/F | - | White | - | - |
| 3 | Pre-TXP | - | 20yo/M | - | White | - | - |
| 4 | Pre-TXP<br>No ACR | N | 15yo/M | 6yo/F | Hispanic | White/NH | PFIC |
| 5 | Pre-TXP<br>No ACR | N | 13yo/M | 14yo/F | White/Hispanic | Black | BA |
| 6 | Pre-TXP<br>Late ACR (n=4) | Y | 10yo/F | 11mo/M | Unknown | Black/NH | Alagille |
| 7 | No ACR<br>Late ACR (n=2) | N | 6mo/M | 8mo/F | White | White/NH | BA |
| 8 | Late ACR | Y | 27yo/M | 18mo/F | White/NH | Black/NH | Hepatoblastoma |
| 9 | Late ACR | Y | 2yo/M | Unknown | Unknown | White/NH | BA |
| 10 | Late ACR | Y | 18yo/M | 10mo/F | Black/NH | White | BA |
| 11 | Late ACR (n=3) | Y | 42yo/M | 14yo/M | White | White/NH | A1AT, HCC |
| 12 | Late ACR (n=2)<br>ACR Resolved (n=3) | Y | 7yo/M | 8mo/F | Unknown | White/NH | BA |
| 13 | Late ACR<br>ACR Resolved | Y | 25yo/M | 8mo/F | White | White/NH | BA |
| 14 | Late ACR<br>ACR Resolved | Y | 15yo/F | 9yo/M | White | Black/NH | BA |

Information on prior ACR, recipient age, sex, race, and ethnicity, and transplant indication are not applicable for Pre-TXP only samples. Abbreviations: A1AT = alpha 1 antitrypsin deficiency, ACR = acute cellular rejection, BA = biliary atresia, GSD = glycogen storage disease, F = female, M = male, NH = non-Hispanic, Pre-TXP = pre-transplant liver, PFIC = progressive familial intrahepatic cholestasis.

**Table 2.** Rejection characteristics.

| Patient | Rejection Category | ALT<br>(units/L) | GGT<br>(units/L) | DSA | Banff<br>score<br>(0-9) | Baseline IS | Tac level<br>(ng/mL) | Rejection Treatment | Post-Transplant<br>Day |
| --- | --- | --- | --- | --- | --- | --- | --- | --- | --- |
| 1 | Pre-TXP | - | - | - | - | - | - | - | - |
| 2 | Pre-TXP | - | - | - | - | - | - | - | - |
| 3 | Pre-TXP | - | - | - | - | - | - | - | - |
| 4 | Pre-TXP<br>BX2: No ACR | -<br>111 | -<br>78 | -<br>Neg | -<br>3 | -<br>TAC+MMF | 10.1<br>- | -<br>- | 69<br>- |
| 5 | Pre-TXP<br>BX1: No ACR | -<br>193 | -<br>33 | -<br>Neg | -<br>ND | -<br>TAC | 8<br>- | -<br>- | 113<br>- |
| 6 | Pre-TXP<br>BX2: Late ACR<br>BX3: Late ACR<br>BX4: Late ACR<br>BX5: Late ACR | -<br>525<br>298<br>234<br>80 | -<br>357<br>954<br>366<br>284 | -<br>Neg<br>Neg<br>Neg<br>Neg | -<br>4<br>3<br>2<br>3 | -<br>TAC<br>-<br>-<br>- | 5.1<br>10.3<br>12.2<br>10.5<br>- | -<br>TAC+Rituximab<br>TAC+Steroids<br>TAC+Steroids<br>TAC+Steroids+MMF | 319<br>331<br>385<br>434<br>- |
| 7 | BX2: No ACR<br>BX3: Late ACR<br>BX4: Late ACR | 31<br>68<br>69 | 214<br>382<br>822 | Neg<br>Neg<br>Neg | 2<br>4<br>5 | TAC<br>-<br>- | 12.2<br>12.1(rapa)<br>3.2 | TAC+Steroids+RAPA<br>TAC+Steroids+RAPA<br>- | 147<br>292<br>306 |
| 8 | BX2: Late ACR | 203 | 46 | Neg | 4 | TAC | 4.3 | TAC+MMF | 477 |
| 9 | BX1: Late ACR | 89 | 254 |  | 4 | TAC | 10.8 | TAC+Steroids | 113 |
| 10 | BX1: Late ACR | 216 | 135 | Pos | 7 | TAC | 11.1 | TAC+Steroids | 1058 |
| 11 | BX1: Late ACR<br>BX2: Late ACR<br>BX3: Late ACR | 278<br>436<br>205 | 165<br>342<br>238 | Pos<br>Neg<br>Neg | 5<br>7<br>5 | TAC<br>TAC<br>- | 9<br>4.9<br>10.3 | TAC+Steroids<br>TAC+Steroids<br>TAC+Steroids+ATG | 1395<br>1772<br>1777 |
| 12 | BX1: Late ACR<br>BX2: Late ACR<br>BX3: Resolved<br>BX4: Resolved<br>BX5: Resolved | 545<br>576<br>149<br>188<br>71 | 309<br>595<br>522<br>353<br>220 | Pos<br>Pos<br>Pos<br>Neg<br>- | 5<br>5<br>0<br>0<br>2 | TAC<br>-<br>-<br>TAC+MMF<br>TAC+MMF | 5.4<br>10<br>10.3<br>11<br>6.2 | TAC+Steroids<br>TAC+Steroids+ATG<br>TAC+MMF<br>-<br>- | 1772<br>1775<br>1820<br>1863<br>2072 |
| 13 | BX1: Late ACR<br>BX2: Resolved | 124<br>347 | 23<br>20 | Neg<br>Neg | 3<br>4 | TAC<br>- | 7.6<br>9.6 | TAC+Steroids<br>- | 237<br>279 |
| 14 | BX1: Late ACR<br>BX4: Resolved | 230<br>295 | 470<br>1122 | Pos<br>Pos | 5<br>3 | TAC+steroids+RAPA<br>- | 8.8<br>11.5 | TAC+Steroids+RAPA+ATG<br>TAC+Steroids+MMF | 3077<br>3119 |

Information on ALT, GGT, DSA, Banff score, baseline IS, Tac level, rejection treatment, and post-transplant day are not applicable for Pre-TXP only samples. Abbreviations: ACR: acute cellular rejection, ATG: anti-thymocyte globulin, BX: biopsy, IS; immunosuppression, MMF: mycophenolate mofetil, ND: not done, Pre-TXP: pre-transplant donor liver, TAC: tacrolimus, RAPA: rapamycin.

After alignment, quality control filtering, sample integration, and dimensionality reduction, we identified 12 broad transcriptionally distinct clusters, which we annotated using known marker genes and cell-type specific gene signatures (**Figure 1A**). These clusters were not isolated to any individual patient or histologic diagnosis, indicating good integration (**Figure 1B**). All cell types were observed and their frequencies relatively consistent across histologic diagnosis (**Figure 1C**). As expected, the proportion of T cell populations (CD4+, CD8+ T cells, and Gamma Delta (γ/δ) T cells) were significantly higher in the Late ACR samples compared to the Pre-Transplant (Pre-TXP) samples (**Figure 1C**). On the other hand, Kupffer cells and Monocyte-Derived Macrophages were significantly less prevalent in Late ACR compared to Pre-TXP (**Figure 1C**).

**Figure 1.**
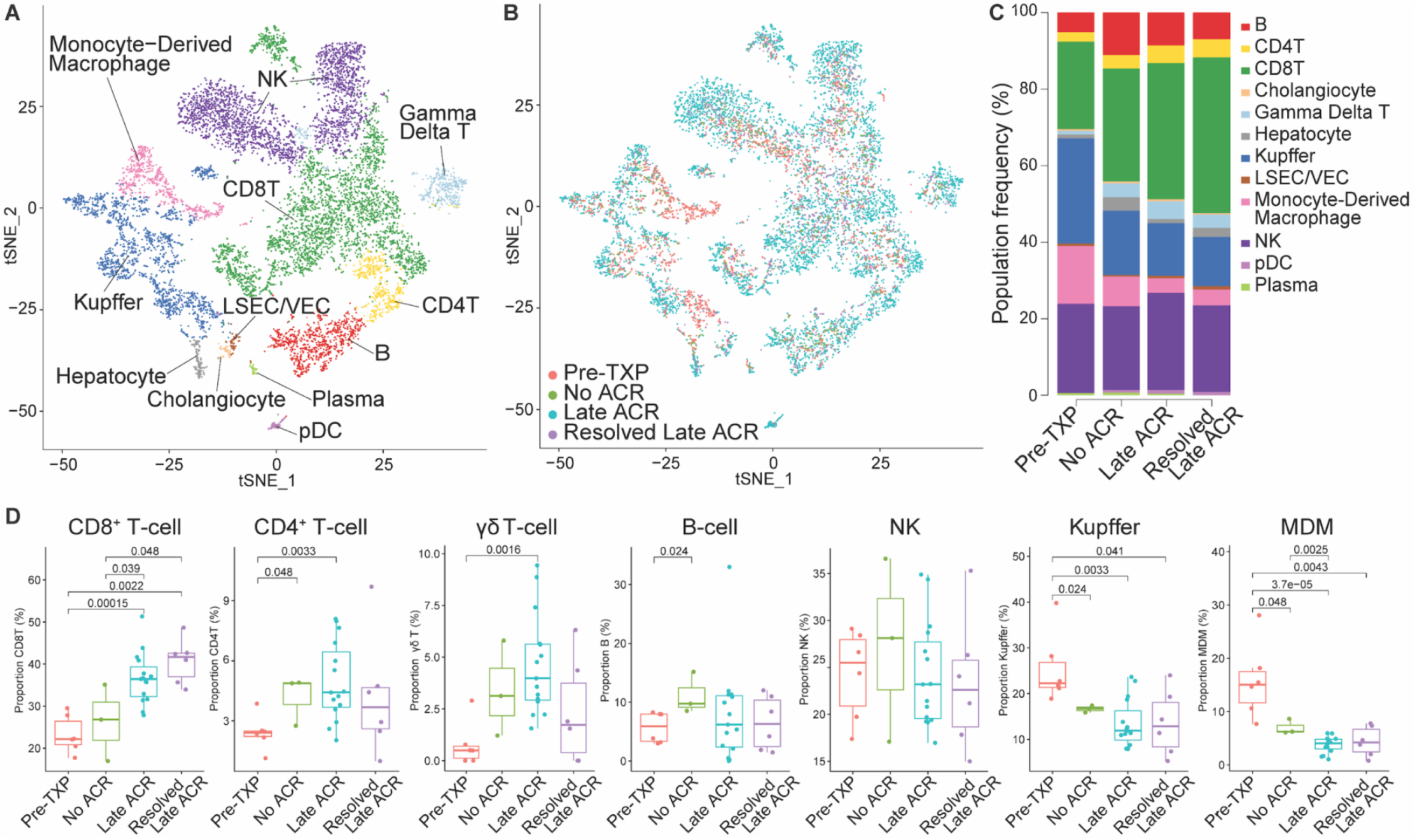
Cell identification for thirty integrated liver biopsy samples. (**A**) TSNE plot showing 10,566 cells from thirty samples across fourteen patients, integrated and clustered using Seurat. Broad cell types are annotated and separated by color. (**B**) TSNE plot colored by histologic diagnosis as indicated in Table 1. (**C**) Stacked bar plots indicating cell type frequencies within each histologic diagnosis, colored by broad cell types. (**D**) Cell population quantification of total recovered cells condition, p-value listed above each cellular comparison. The unlabeled columns have a p-value >0.05 (not significant). Abbreviations: Acute cellular rejection (ACR), B-cell (B), CD4^+^ T cell (CD4T), CD8^+^ T cell (CD8T), liver sinusoidal endothelial cell (LSEC), natural killer cell (NK), plasmacytoid dendritic cell (pDC), vascular endothelial cell (VEC).

### Expanded CD8^+^ T cell clones persist in serial biopsy samples despite treatment

To gain more clarity into the T cell subpopulations present in ACR, the T cell populations were re-integrated and clustered at a higher resolution, resulting in 7 distinct T cell subpopulations (**Figure 2A**). These T cell populations were annotated using a combination of known T cell markers starting with their expression of *CD3γ*, *CD4* or *CD8ɑ*, and either *TRA* or *TRD*. The proportions of T cell subclusters were consistent across histologic diagnoses with the exception of γ/δ T cells, which were significantly increased in Late ACR relative to Pre-TXP, and significantly decreased in Resolved Late ACR relative to Late ACR (**Figure 2B**). Individual T cell subtypes were annotated as follows: CD4^+^ naive T cells (CD8_NAIVE_) were identified by their high expression of *CD4, CCR7,* and *SELL*; MAIT cells were identified by expression of *TRAV1-2* (*50*); γ/δ T cells were identified by their high expression of TRDV1 along with a lack of detected *TRA* and *TRB* chains in the V(D)J analysis. CD8^+^ Effector T (CD8_EFF_) cells were identified by high expression of *CD27, PRF1, GZMB,* and *CD69* with concomitant decreased expression of the tissue egress molecule S1PR1. CD8^+^ Effector Memory T cells (CD8_EM_) were distinguished by low/absent CD27, increased S1PR1 expression, and persistently high expression of the effector molecules *PRF1* and *GZMB* (51, 52). Recently, a CD56^+^CD161^-^ NK-like CD8^+^ T cell subpopulation in the liver was described, which is present in the sinusoids, bears some tissue residence markers including CD69 and CD103, has a restricted TCR repertoire, and is characterized by increased expression of NK markers including *NCAM1* and *KLRC2* (47). We uncovered a very similar population in our intragraft single cell analysis, identifying a subpopulation of CD8^+^ T cells expressing *NCAM1, KLRC1/C2*, and *KLRD1* but lacking expression of *Eomes* and *CD161/KLRB1* (**Figure 2C**). Because of their similarity to the previously-described CD56^+^CD161^-^ CD8^+^ T cell subpopulation, we termed these cells ‘CD8^+^ NK-like T cells (CD8_NK-like_).’ Finally, the Cycling T cluster was defined by high G2/M-phase and S-phase scores derived with Seurat as well as their high expression of *MKI67* (**Figure 2D**).

**Figure 2.**
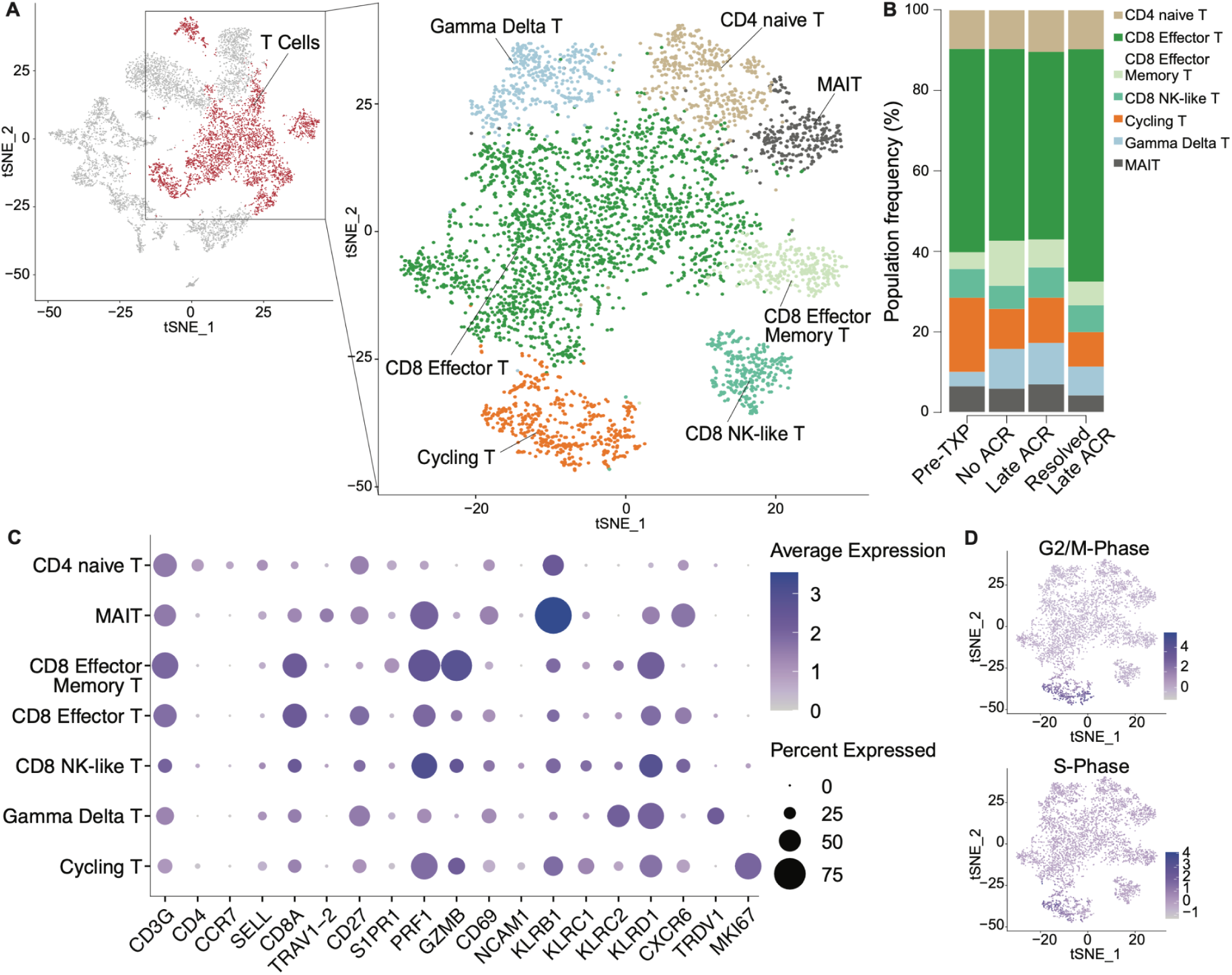
T cell subtype identification. (**A**) CD4^+^ T cells, CD8^+^ T cells, and gamma delta T cells were isolated and clustered separately to define additional subpopulations of cells and visualized on a TSNE plot. (**B**) Stacked bar plots indicating T cell subtype frequency for each histologic diagnosis. γ/δ T cells were significantly increased in Late ACR vs. Pre-TXP, and decreased in Late ACR vs. Resolved Late ACR (p<0.05). The remaining T cell subcluster frequencies did not achieve statistical significance amongst histologic diagnoses. (**C**) Dot plot showing average expression for key T cell defining genes (indicated by color) and the percent of cells within each T cell subtype that express that gene (indicated by dot size). (**D**) TSNE plot with cell cycle phase scoring, with darker blue indicating a higher score. Abbreviations: Acute cellular rejection (ACR), mucosal associated invariant T cell (MAIT).

We next examined the intragraft TCR gene expression for evidence of T cell clonal expansion. First, the *TRA* and *TRB* CDR3 V(D)J sequence outputs from Cell Ranger were matched to the corresponding barcoded cells within our gene expression dataset. Of the cells with detectable *TRA* or *TRB*, 85.05% expressed both *TRA* and *TRB* sequences concurrently, while only 2.76% expressed isolated *TRA*, 12.18% expressed isolated *TRB* (**Supplemental Figure 1**). T cells that shared the identical *TRA* and *TRB* sequences were considered to have the same clonotype, and clone size was quantified for each clonotype based on the number of cells that shared that clonotype. Further, clonal expansion was defined as any clonotype with a clone size of 3 or greater, ensuring the assessment of cells that had divided at least twice. Consistent with their gene expression, the CD4^+^ naive T cells do not contain any expanded TCR sequences and the vast majority of cells in this cluster contain unique, unexpanded *TRA/B* pairs (**Figure 3A, B**). Because γ/δ T cell subpopulations do not express *TRA* or *TRB*, CD8_EXP_ did not reside within this subset (**Figure 3A, B**). Rather, the cell types containing the highest frequency of clonally expanded TCR fell within the CD8_EFF_ subcluster with some CD8_EXP_ clones found in the cycling, CD8_EM_, and CD8_NK-like_ subclusters (**Figure 3A-C**). Of the CD8_EXP_ clones, only ∼10% contained dual alpha chains (*TRA*/*TRA*/*TRB*), which was not significantly different from the frequency of dual-*TRA* usage in CD8_UNEXP_ clones (7.6%). The number of CD8_EXP_ clones were significantly increased in late ACR biopsies compared to those found in pre-transplant control liver biopsies (Pre-TXP), with a trend toward decreased CD8_EXP_ in both resolved and recurrent ACR (**Figure 3D**, upper right panel). Consistent with their presumed role in alloreactivity, the number of cells per each CD8_EXP_ clone showed an increased trend in Late ACR as compared to either Pre-TXP donor liver or No ACR biopsies, and significantly decreased in Resolved ACR (**Figure 3D**, lower right panel).

**Figure 3.**
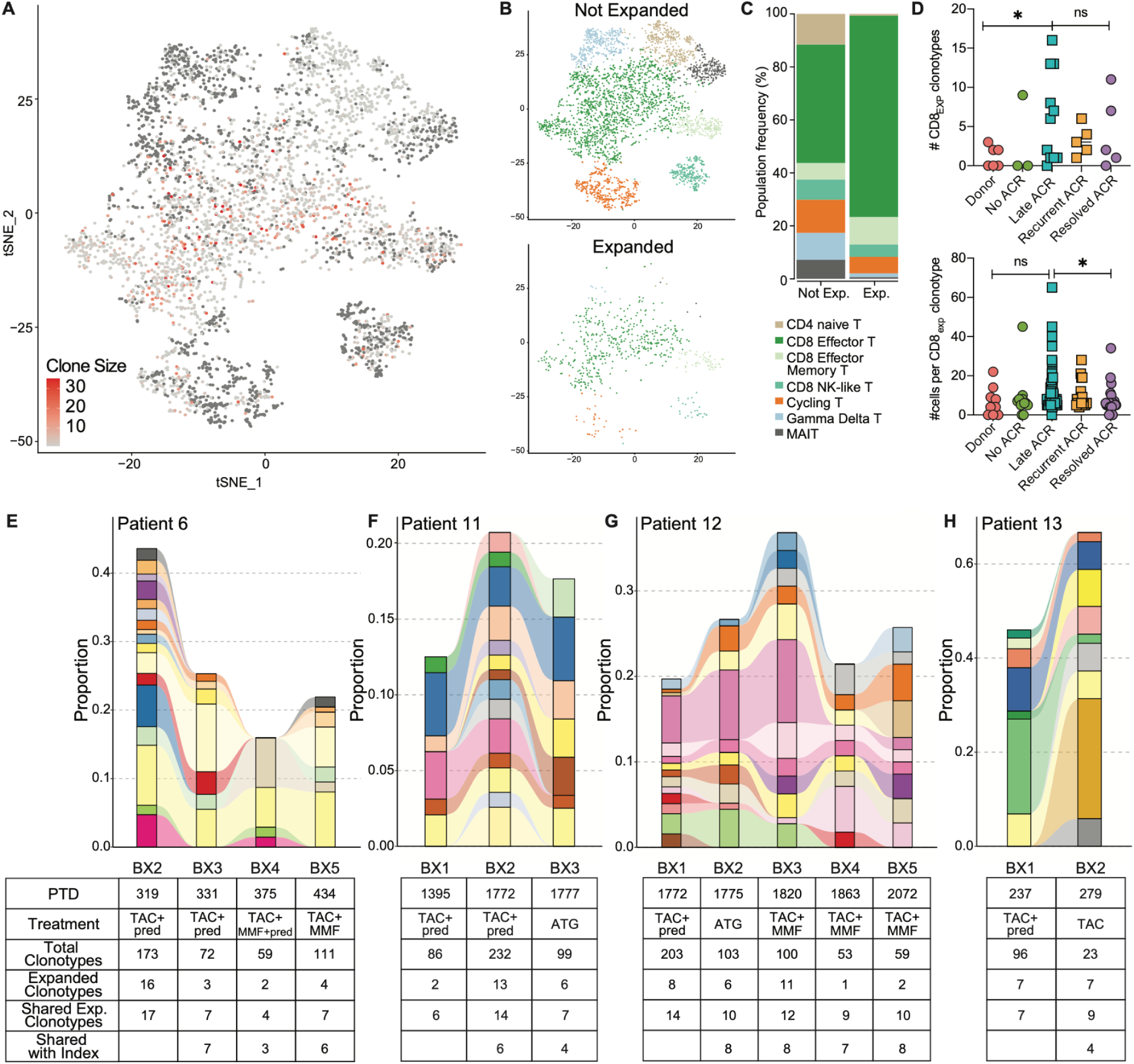
CD8^+^ expanded T cell receptor clones persist in serial biopsies despite anti-rejection treatment. (**A**) TSNE plot of T cells with superimposed clone size. red=high clone size, light gray=no clonal expansion, dark gray=no TRA/B detected. (**B, C**) Distribution of CD8_EXP_ intragraft clonotypes amongst T cell subpopulations. (**D**) Number of expanded CD8 clonotypes (CD8_EXP_) and number of cells per CD8_EXP_ clonotype were quantified, *=p<0.05, ns=not significant. (**E-H**) Connected stacked bar plots for Patients 6, 11, 12, and 13 showing shared CD8_EXP_ clones across biopsies. The y-axis indicates the proportion of T cells containing each clonotype within a sample, with distinct clonotypes marked by separate colors. Swaths of color between two side-by-side bar plots indicate clonal detection in each biopsy. The tables below each bar plot describe the post-transplant day (PTD) for that sample, anti-rejection treatment initiated after each biopsy, the number of distinct clonotypes, and the number of clonotypes shared with the first (index). Abbreviations: Acute cellular rejection (ACR), Antithymocyte globulin (ATG), expanded (exp), Mycophenolate mofetil (MMF), prednisone (pred), Tacrolimus (TAC).

Because shared CD8_EXP_ clonotypes were detected in serial biopsies from each individual patient, we next determined whether these CD8_EXP_ clonotypes were shared between patients, thereby representing potential ‘public’ allospecific clonotypes. Of note, neither the patients nor their donors had any common HLA alleles. Of the 99 shared expanded intragraft *TRA/B* clonotypes detected in rejection biopsies, none were detected in either pre-transplant donor liver or No ACR biopsies (**Supplementary Figure 2**). However, only eight of the shared intragraft CD8_EXP_ clonotypes were shared between two patients (patient 11 and patient 12), neither of whom had any HLA alleles in common from either the donor or recipient standpoint (**Supplementary Figure 2**). Analysis of these eight shared CDR3 sequences in the public TCR database, vdjdb.cdr3.net, did not yield any previously reported epitope specificity. Taken together, these results suggest that the CD8_EXP_ clonotypes are alloreactive and represent largely ‘private’ specificities.

We next determined, in individual patients (patients 6, 11, 12, and 13), whether the CD8_EXP_ clonotypes that were present at the time of rejection were also present in subsequent biopsies. We generated connected stacked bar plots to visualize the proportion of expanded T cells within each of the shared clonotypes (**Figure 3E-H**). We found that multiple large CD8_EXP_ clones remained detectable and expanded in subsequent liver biopsies from the same patient (**Figure 3E-H**). Surprisingly, the majority of CD8_EXP_ clones detected at index rejection in all 4 patients were retained in the liver of patients up to 377 days between rejection episodes (as in the case of patient 11) (**Figure 3F**). In two patients with recurrent rejection (patients 6 and 11), the vast majority of the CD8_EXP_ intragraft clonotypes remained detectable in all subsequent biopsies despite each patient receiving increased immunosuppression with tacrolimus, corticosteroids and/or mycophenolate mofetil (MMF). In the case of patient 12, identical CD8_EXP_ intragraft clonotypes were detected despite administration of anti-thymocyte globulin (ATG) (**Figure 3G**). Although ATG effectively depleted the peripheral lymphocyte compartment in patient 12 from 52% to 14% CD3^+^ and 1004 to 161 CD8^+^ cells/µL, intragraft CD8_EXP_ clonotypes remained detectable (**Figure 3G**). In two patients with histologically resolved rejection (patients 12 and 13), identical CD8_EXP_ clonotypes were consistently detected in the liver allograft despite histologic resolution of the rejection episode (**Figure 3G, H**). Taken together, these results suggest that CD8_EXP_ intragraft clonotypes remain detectable in serial biopsies from rejecting LTx patients despite augmented immunosuppression and treatment with a T-depleting polyclonal antibody preparation (ATG).

Although the scRNAseq data allowed identification of CD8_EXP_ clonotypes, these data did not inform their location within the rejecting allograft. To determine the spatial distribution of CD8_EXP_ clonotypes during active and resolved rejection, we designed RNA probes specific for the most abundant TCRɑ/β clone found in patient 6 and a separate clone in patient 12. Notably, neither these patients nor their donors have any HLA alleles in common. Using these probes and the BaseScope dual index assay on FFPE liver tissue, we found that the CD8_EXP_ clone from patient 6 (who had multiple episodes of recurrent ACR) was detected within inflamed portal areas at all biopsy timepoints, in close proximity to injured bile ducts (**Figure 4**, arrows). For patient 12, the CD8_EXP_ clone was detected initially in inflamed portal areas at index rejection, but after histologic resolution was detected only in lobular infiltrates distant from bile ducts, remaining detectable even after treatment with ATG (**Figure 5**, arrows). These data validate the scRNAseq findings and suggest that although intragraft CD8_EXP_ clones remain detectable within the allograft after rejection resolves, they lack the ability to traffic to portal areas.

**Figure 4.**
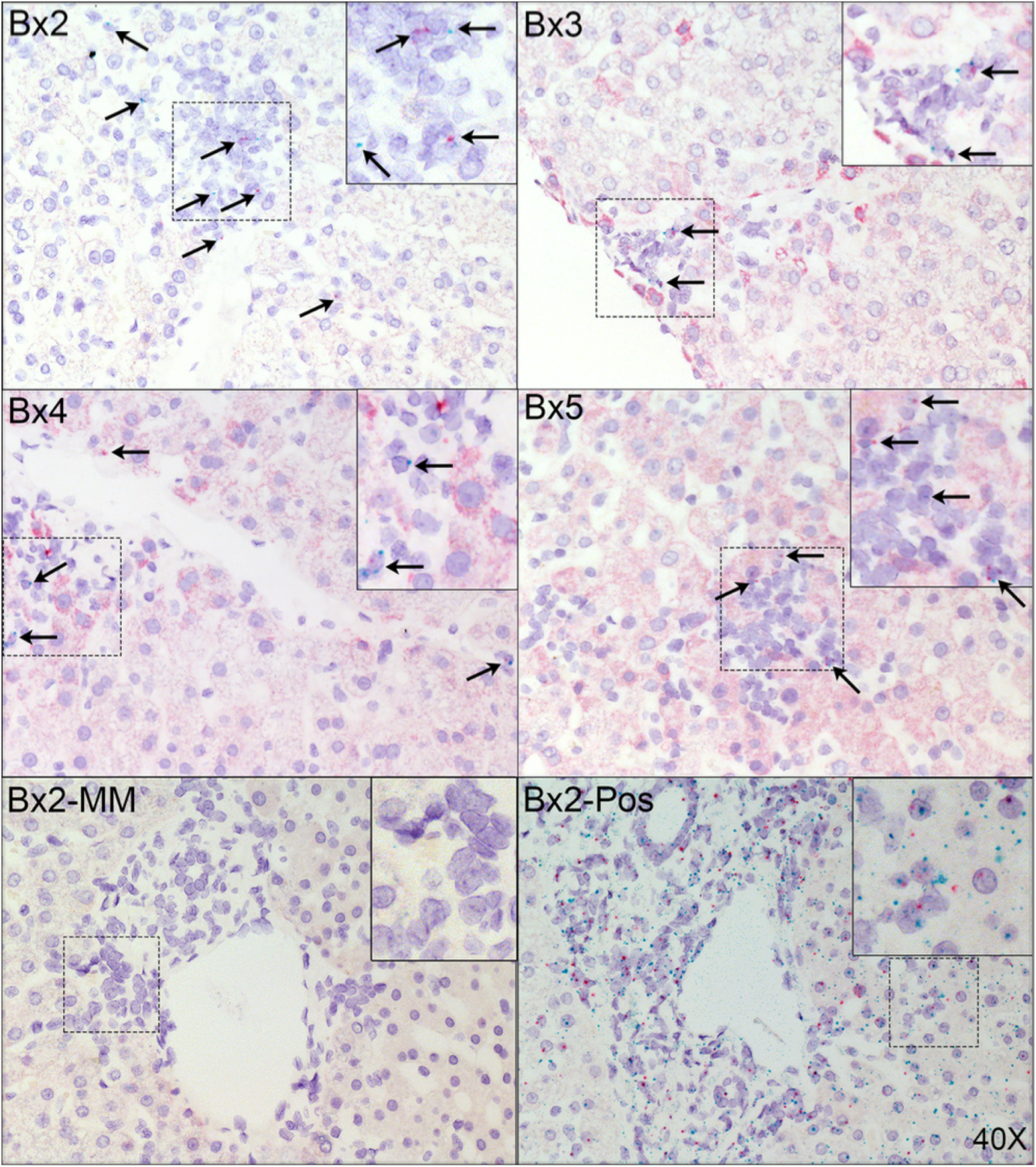
CD8_EXP_ TCR clones are detected primarily in inflamed portal tracts of liver biopsy tissue obtained from a patient with ongoing late ACR. BX2-5 are shown from Patient 6. Black arrows highlight CD8_EXP_ clones. Images obtained using brightfield microscopy at 40x magnification. Probe control (Mismatch) from Patient 12 is shown in the lower left panel, while commercial positive control probes for housekeeping genes is shown in the lower right. Dashed rectangle highlights inflamed portal tracts and is magnified and shown in the upper right inset of each panel.

**Figure 5.**
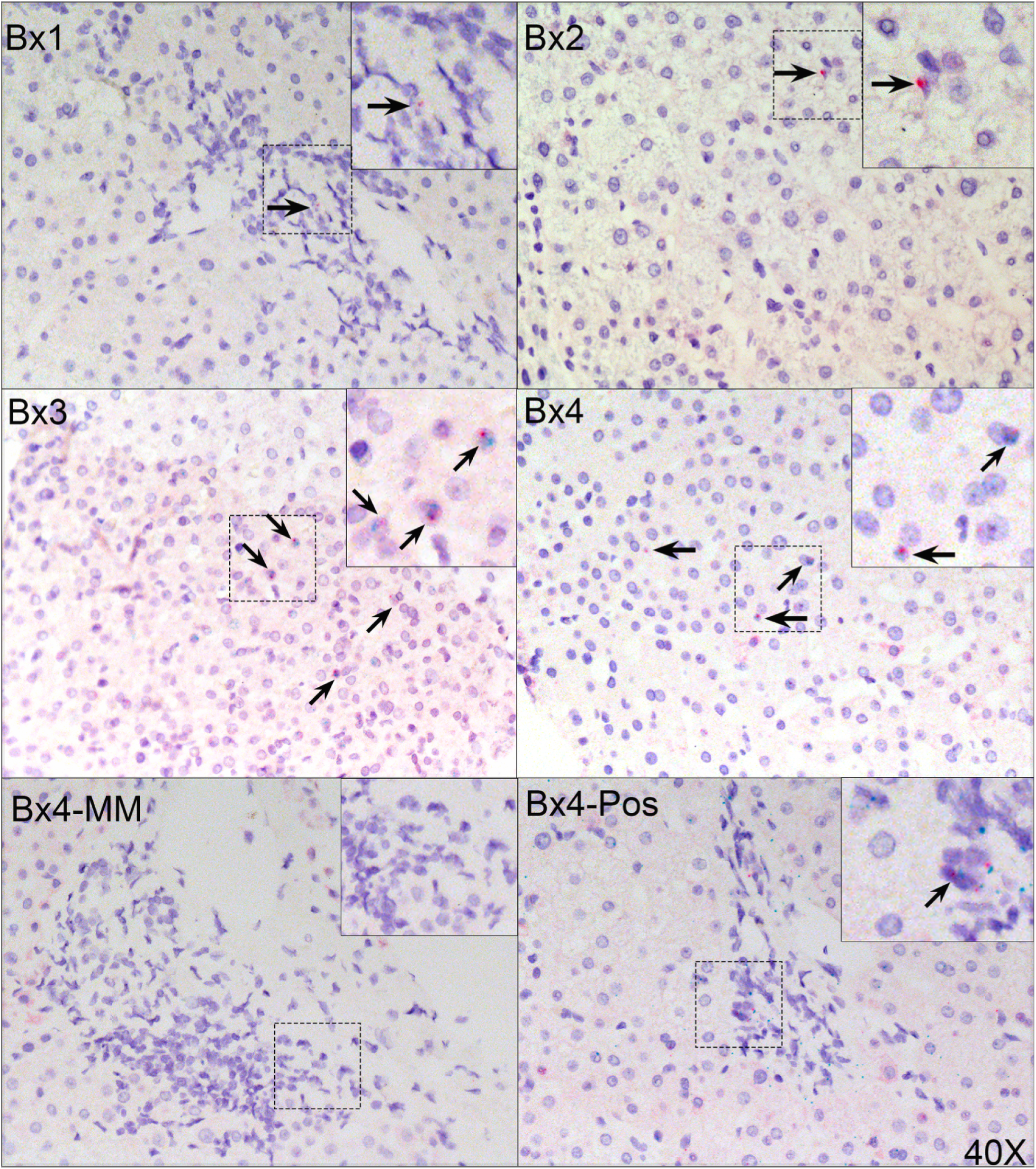
CD8_EXP_ TCR clones are detected primarily in lobular areas of liver biopsy tissue obtained from a patient with histologically resolved late ACR. BX1-4 are shown from Patient 12. Black arrows highlight CD8_EXP_ clones with punctate red and green staining. Probe control (Mismatch) from Patient 6 is shown in the lower left panel, while commercial positive control probes for housekeeping genes shown in the lower right. Images obtained using brightfield microscopy at 40x magnification. Dashed rectangle highlights lobular inflammation and is magnified and shown in the inset of each panel.

Although the CD8_EXP_ T cell clones largely remained within the CD8_EFF_ subcluster, we next addressed whether they changed phenotype over time by performing supervised gene expression analysis in CD8_EXP_ vs. CD8_UNEXP_ T cell clones, stratified by disease state. Level of expression of genes associated with activation (*GZMB, HLA-DRA*), exhaustion (*LAG3, TIGIT, NKG7*), and tissue residence (*CD69, CXCR6*) were measured in CD8_EXP_ clones relative to CD8_UNEXP_ clones in the four patients with serial biopsies (patients 6, 11, 12, and 13). As expected, violin plots show upregulation of *GZMB* and *HLA-DRA* expression, as well as CXCR6 expression, during index rejection (**Figure 6**). As expected, in the biopsies from patients with recurrent rejection, activation markers persisted without evidence of developing exhaustion (**Figure 6**). Surprisingly, in patients with resolved rejection, CD8_EXP_ did not have evidence of exhaustion but instead retained high expression of activation and tissue residence markers. These data, combined with those from **Figures 4** and **5** demonstrating continued lobule-restricted presence of CD8_EXP_ clones once rejection resolves, suggest that intragraft CD8_EXP_ remain active but lack the ability to traffic to portal areas once the rejection episode is resolved.

**Figure 6.**
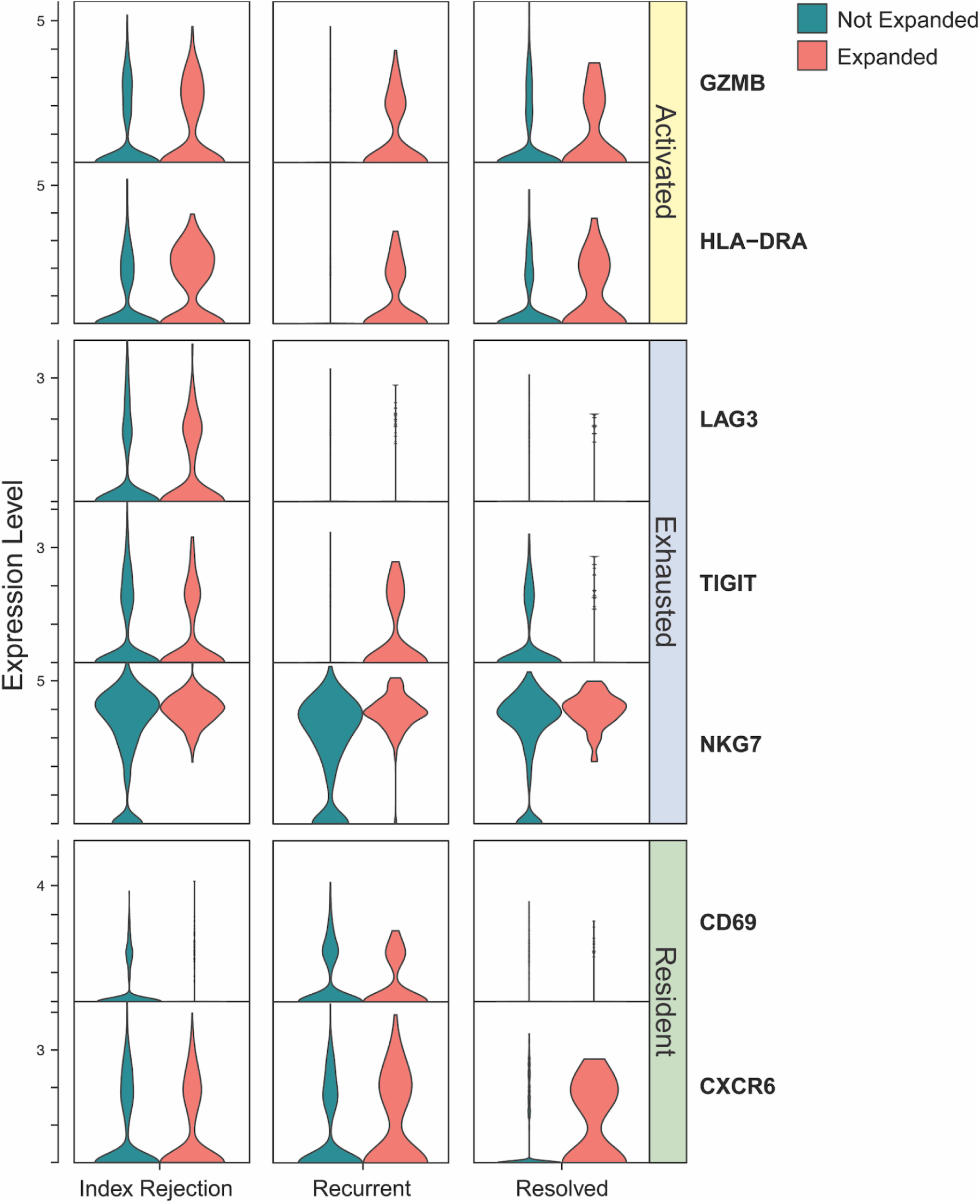
CD8_EXP_ TCR clones retain activation and tissue residence markers despite histologic resolution of rejection. Supervised gene expression analysis of activation, exhaustion, and tissue residence markers in CD8_UNEXP_ T cell clones (green violin plots) vs. CD8_EXP_ (red violin plots) of patients with serial biopsies (patient 6, 11, 12, 13). CD103 was not detected.

To determine whether B cells were also clonally expanded and potentially assisting T cells in forming tertiary lymphoid structures in the allograft, we utilized the V(D)J data within the same dataset to track B cell clonal expansion across histologic diagnoses. The B-cell population was identified by expression of *MS4A1* (CD20) and the presence of immunoglobulin transcripts. Further sub-clustering identified two clusters: Naive B-cells and Transitional B-cells (**Supplemental Figure 3A**), with Transitional B-cells expressing higher levels of *CD24* and *CD38* (**Supplemental Figure 3C**). Cell type proportions for B-cell subclusters were not significantly different across histologic diagnoses (**Supplemental Figure 3B**). We next quantified clone size for B-cells and found that the majority of B-cells were not clonally expanded, which was independently validated by a separate B-cell pipeline (**Supplemental Figure 3D**). Although a high level of somatic hypermutation could potentially mask true expanded clones within the dataset, most of the B-cells had relatively low levels of hypermutation (**Supplemental Figure 3E**). Taken together, the intragraft B cell subsets were primarily of naive and transitional phenotype and lacked evidence of somatic hypermutation or clonal expansion. This finding is in keeping with the low level of DSA present in the majority of the patients in our dataset and the lack of histologically identified features of antibody-mediated rejection.

### NK subpopulations do not correlate with rejection activity

NK cells are a rich source of cytokines, including IL-15, which can activate and preserve tissue-resident CD8^+^ T cells, and are critical for control of CMV infection in liver transplant recipients (53, 54). We next investigated whether NK cell subsets differed between histologic rejection subtypes. The NK cell population was subset into two distinct clusters: CD56^bright^ and CD56^dim^ (**Supplemental Figure 4A**). There were no significant differences in the prevalence of these subpopulations across histologic diagnoses (**Supplemental Figure 4B**). CD56^bright^ cells show higher expression of *CD56* (*NCAM1*), *IL2RB*, *EOMES*, and *CD160*, while cells in the CD56^dim^ population have higher expression of *CX3CR1* and *GZMB* (**Supplemental Figure 4C**).

### Kupffer cell subpopulations differ in composition in late ACRs

The liver contains a diverse array of tissue-resident and infiltrating macrophage and Kupffer cell (KC) subsets which are vital to stimulation of T cell subpopulations within the liver (55, 56). To determine whether there were differences in macrophage and KC populations in rejecting and non-rejecting liver tissue, we next performed high resolution sub-clustering of KC and monocyte-derived macrophages. This resulted in 5 transcriptionally distinct KC/macrophage clusters (LST1^+^, CD1C^+^, C1Q^+^, PTPRC^+^, and IFI27^+^ subclusters, respectively) and the original Monocyte-Derived Macrophage cluster (**Figure 7A**). Across histologic diagnosis, we observed statistically significant differences in the proportion of cells in each cluster (**Figure 7B**). The CD1C^+^, LST1^+^, and PTPRC^+^ KC populations showed significant increases in Late ACR samples compared to Pre-TXP, while the IFI27^+^ KC and Monocyte-Derived Macrophages showed a significant decrease.

**Figure 7.**
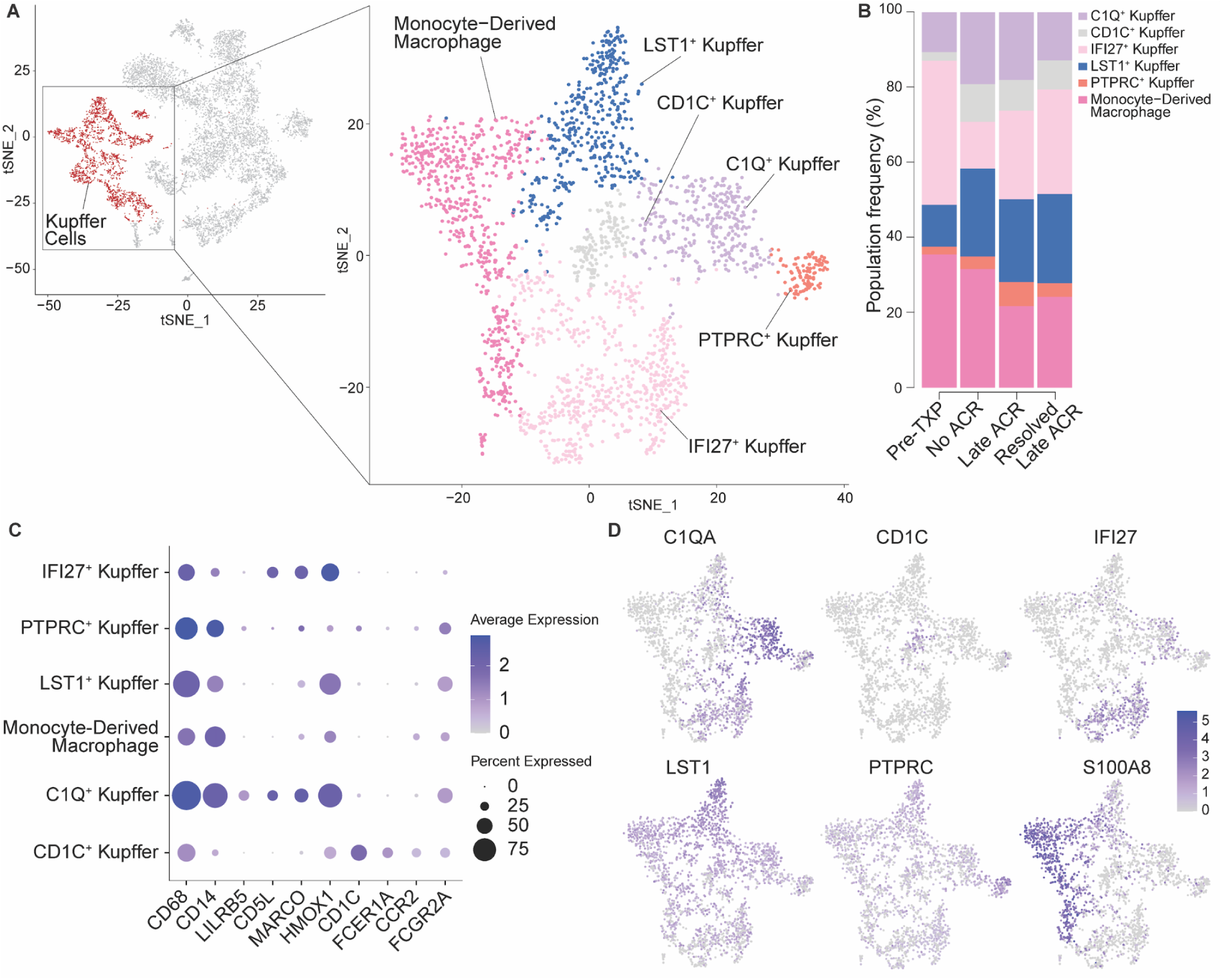
Kupffer cell subtype identification. (**A**) Kupffer cells and monocyte-derived macrophage populations were subset from the larger dataset and reclustered to identify unique subpopulations and visualized with TSNE. (**B**) Stacked bar plot indicating subtype frequency with respect to histologic diagnosis. The CD1C+, LST1+, and PTPRC+ populations were significantly increased, while IFI27+ and monocyte-derived macrophages were significantly decreased in Late ACR vs. Pre-TXP (p<0.05). C1Q+, LST1+, and CD1C+ populations were significantly increased in No ACR vs. Pre-TXP (p<0.05). LST1+ and CD1C+ populations were significantly decreased in Resolved Late ACR vs. Pre-TXP (p<0.05). The remaining subset comparisons were not statistically significant. (**C**) Dot plot showing average expression for key Kupffer cell defining genes (indicated by color) and the percent of cells within each Kupffer cell subtype that express that gene (indicated by dot size). (**D**) TSNE plots with cells colored by expression of each of the genes used to determine names for the unique subpopulations (*C1QA, CD1C, IFI27, LST1,* and *PTPRC*) as well as *S100A8* to clearly define the monocyte-derived macrophage population. Dark blue indicates higher expression of each gene and light gray indicates lower expression on a shared scale. Abbreviations: Acute cellular rejection (ACR).

Further, C1Q^+^ KC were more abundant in No ACR versus Pre-TXP (p = 0.024). We also observed that both LST1^+^ and CD1C^+^ KC were significantly more abundant in No ACR and Resolved Late ACR compared to Pre-TXP. However, there are no significant differences between KC/macrophage subsets in late ACR vs. No ACR, suggesting that differences between Late ACR and Pre-TXP samples are due to immunosuppression and/or the transplant process rather than specific changes related to rejection. Expression of key cell type defining genes were used to identify the Kupffer population (**Figure 7C**) and differential expression analysis was used to label the unique Kupffer cell subclusters and Monocyte-Derived Macrophages (**Figure 7D**).

### Differential crosstalk between KC/macrophage subsets and CD8_EXP_ T cell clones

An increased frequency of APC:CD8^+^ T contacts via immune synapses in hepatic lobules are associated with rejection during immunosuppression withdrawal in pediatric liver transplant recipients (18–21). Because we strongly suspect that alloreactivity is enriched in CD8_EXP_ cells, we compared the composite expression of a 200-gene rejection associated gene set (derived from the immunosuppression withdrawal trials: iWITH) in our scRNAseq dataset (**Supplemental Figure 5**). The expression of the iWITH rejection-associated gene signatures is overrepresented in CD8_EFF_ and CD8_NK-like_ T cells subsets within our scRNAseq dataset, suggesting that the CD8:APC T cell pairs identified in immunosuppression withdrawal trials may be CD8_EXP_ (**Supplemental Figure 5**). We next performed CellPhoneDB analysis to delineate receptor-ligand interactions between KC/macrophage populations and CD8_EXP_ T cells, relative to their CD8_UNEXP_ (bystander) counterparts (**Figure 8A**).

**Figure 8.**
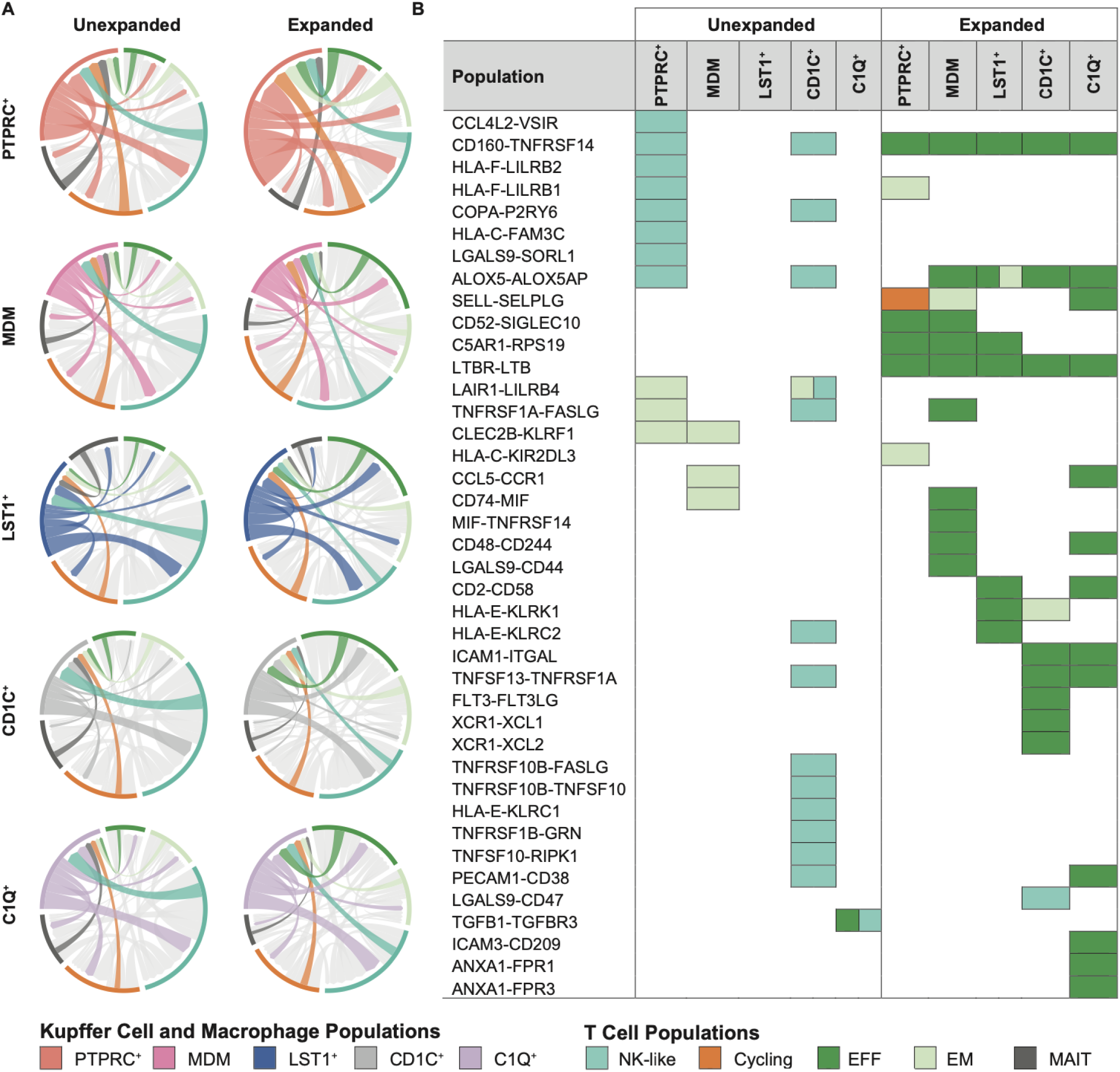
Differential macrophage-T cell crosstalk between macrophage subpopulations and CD8_EXP_ T cell populations. (**A**) CellphoneDB analysis of predicted receptor-ligand interactions between the 5 major macrophage and Kupffer cell (KC) populations and CD8_EXP_ T cells as compared to CD8_UNEXP_ cells. Grayed out interactions are between multiple T cell populations. (**B**) Diagram depicting individual unique receptor-ligand interactions between macrophage/KC subpopulations and CD8_EXP_ vs. CD8_UNEXP_ T cell populations. CD160-TNFRSF14, LTBR-LTB, and ALOX5-ALOX5AP are specifically detected in KC/macrophage interactions with expanded CD8_EFF_.

IFI27^+^ KC did not have significant interactions with either CD8_EXP_ or CD8_UNEXP_ T cells, so was excluded from further analysis. However, the remaining five KC/macrophage subpopulations had increased interactions with expanded CD8_EFF_ (**Figure 8A**, right panel). The CellphoneDB outputs were curated to show only those uniquely detected between a KC/macrophage subpopulation and CD8_UNEXP_ or CD8_EXP_ T cells (**Figure 8B**). Surprisingly, there were significant unique interactions of PTPRC^+^ KC/Macrophage subpopulations or CD1c^+^ KC/Macrophage subpopulations with unexpanded CD8_NK-like_ which were largely inhibitory (*LILRB1/2, LGALS9, FASLG, TGFB1-TGFBR3*; **Figure 8B**). Similarly, interactions between CD8_EXP_ NK-like T cells and the CD1C^+^ KC/macrophage subpopulation uniquely identified *LGALS9-CD47*, an immune checkpoint inhibitor, as a predicted receptor-ligand pair (57–59). In contrast, interactions between KC/Macrophage subpopulations and expanded CD8_EFF_ were primarily activating, with *CD160-TNFRSF14* (LIGHT)*, ALOX5-ALOX5AP* (lipoxygenase pathway), and *LTB-LTBR* receptor-ligand interactions occurred specifically between all five KC/macrophage subpopulations and CD8_EXP_ T cells, but not between KC/macrophage subpopulations and CD8_UNEXP_ T cells (**Figure 8B**). The LTB-LTBR interaction is of particular interest, as this pathway is strongly detected in total liver RNAseq analysis of pediatric LTx recipients who fail to wean from immunosuppression (18) and is required for peripheral lymphoid organ development (60). We attempted to quantify LTB protein by immunofluorescence in rejecting liver tissue. Although LTB^+^CD68^+^ macrophages were detected primarily within the sinusoids and liver lobule, the high background staining of LTB in hepatocytes prevented automated image analysis and quantification. Further validation studies of KC/macrophage activating and inhibitory signaling pathways in rejecting vs. non-rejecting allograft liver tissue are needed in order to understand how intragraft CD8_EXP_ T cell subpopulations are retained and regulated in the liver.

## Discussion

Ours is the first study to successfully perform TCR V(D)J sequencing in combination with single cell RNA sequencing to identify clonally expanded intragraft T cells and transcriptionally define their phenotypes during rejection. Not only were these clones detected at index rejection, but they persisted in subsequent for-cause liver biopsies up to nearly 1 year post ACR treatment, remained significantly expanded in subsequent rejection biopsies, and were localized to both portal and lobular areas of inflammation. Consistent with persistent CD8_EFF_ activation (61), intragraft CD8_EXP_ T cell clones maintained expression of *HLA-DRA, GZMB*, and *CXCR6* in histologically resolved biopsies without developing clear evidence of exhaustion with only minimal upregulation of *PDCD-1*, *LAG3*, and *TIGIT*, and lack of expression of Tox, the major transcriptional regulator of exhaustion (31, 62). In our liver scRNAseq, CD4^+^ T cells and B cell subpopulations, unlike CD8^+^ T cells, were not clonally expanded and primarily of a naive phenotype. Using the bioinformatics receptor-ligand prediction tool CellPhoneDB, we were able to identify unique interactions between KC/macrophage subtypes and CD8_EXP_ T cells, particularly of the CD8_EFF_ phenotype, which identifies candidate pathways which may be important for CD8_EXP_ T cell retention in the rejecting liver allograft.

Our finding that CD8_EXP_ T cells are found in the liver lobule corroborates prior evidence that pediatric late ACR has histologically distinct areas of lobular inflammation (16). Furthermore, prior studies on pediatric LTx who underwent medically supervised immunosuppression withdrawal demonstrated that increased CD8^+^ T cell infiltrates within the hepatic lobule and increased CD8-APC contacts in lobular areas predicted rejection during the withdrawal period and failure of operation tolerance (18–21). Comparison of the iWITH trial liver biopsy rejection-associated transcriptome to genes within CD8_EXP_ T cell clones and with KC subpopulations shows significant overlap, suggesting that the CD8:APC T cell pairs identified in immunosuppression withdrawal trials may be CD8_EXP_ (**Supplemental Figure 5**). Analysis of predicted receptor-ligand interactions between KC/Macrophage subpopulations and unexpanded or expanded CD8 T cell subpopulations yielded a mix of activating and inhibitory interactions. Interactions between PTPRC^+^ KC/Macrophage subpopulations or CD1c^+^ KC/Macrophage subpopulations with unexpanded and unexpanded CD8_NK-like_ T cell subpopulations which were largely inhibitory (*LILRB1/2, LGALS9, FASLG, TGFB1-TGFBR3, LGALS9-CD47*; **Figure 8B**). These inhibitory interactions may serve to constrain activation and proliferation of CD8_NK-like_ T cells in the liver allograft, as CD47 is an immune checkpoint inhibitor which delivers a “don’t eat me” signal to macrophages (57) and is required for acceptance of hepatocyte transplants in the mouse (58, 59). In contrast, interactions between KC/Macrophage subpopulations and expanded CD8_EFF_ were primarily activating. Further understanding of intragraft CD8_EXP_ T cell clone trafficking, activation, and crosstalk with APC is sorely needed, as this knowledge may aid in increasing the frequency of successful pediatric immunosuppression withdrawal to >37.5% of highly selected recipients (19).

Prior studies on TCR sequencing have been limited primarily to the peripheral blood, showing that MLR-reactive CD8^+^ T cell clones are less diverse post-LTx (36, 63–65). While the authors hypothesize that these donor-reactive clones are deleted, the possibility exists that these donor-reactive clones are instead confined to the allograft post-LTx (63). Indeed, Mederacke et al. to date is the only study to directly compare peripheral blood T cell immune repertoire (via Vβ sequencing) to that of the stable and rejecting allograft (36). Although the CD8+ T cell repertoire is less diverse post-LTx, there is no overlap of the peripheral blood compartment in stable LTx recipients or those with subclinical rejection, and only a minority (∼16%) of CD8^+^ T cell clones were shared between the blood and liver during ACR (36). The persistent upregulation of CXCR6 (a marker of tissue residence in the liver) on CD8_EXP_ T cell clones in histologically resolved rejection suggests the possibility that the intragraft CD8_EXP_ T cell clones are taking on characteristics of tissue residence and may instead be confined to the allograft. While we cannot currently exclude the possibility that CD8_EXP_ clones are recirculating from peripheral blood at each biopsy timepoint, we are actively investigating the potential clonal overlap between peripheral alloreactive CD8_EXP_ T cell clones and those in rejecting liver tissue.

Our analysis of the shared CD8_EXP_ *TRA/B* sequences shows little clonal overlap between patients, confirming that alloreactivity relies largely on private specificities. However, the specificities of CD8_EXP_ clonotypes remains unclear. It is possible that CD8_EXP_ clonotypes may represent pre-existing memory cells specific to viral peptides in the context of recipient HLA, or instead recognize donor HLA molecules with unknown peptides (51, 66, 67). A third possibility is that CD8_EXP_ clonotypes are cross-reactive, recognizing both recipient HLA:viral peptide ligands and simultaneously recognizing donor HLA peptides in the context of donor HLA presentation, as recently demonstrated in a mouse heart transplantation model (68). Although prior studies on kidney allograft rejection defined the intragraft CD8_EXP_ as alloreactive (37), it remains possible that some of the CD8_UNEXP_ T cell clones within our dataset are either directly alloreactive or contribute to the alloresponse by providing costimulatory ‘help’ to alloreactive CD8_EXP_ clones, as recently shown in a mouse skin transplantation model (69). The avidity, allospecificity, viral peptide cross-reactivity, and potential cooperation between CD8_EXP_ vs. CD8_UNEXP_ T cell clones is a subject of ongoing investigation utilizing human CDR3-expressing Jurkat 76 T cell lines recently described by our group (37).

Limitations of our study include the relatively low representation of hepatocytes, cholangiocytes, and endothelial cells within our dataset. This technical challenge is common to others seeking to understand the immune compartment of the liver (70), but unfortunately precludes full analysis of T cell-target cell crosstalk and identification of cell injury patterns in endothelial cells, cholangiocytes, and hepatocytes, all of which are known cellular targets for allorecognition during LTx rejection. However, we are adapting single nuclear RNA sequencing protocols to cryopreserved liver biopsy specimens which provides the full repertoire of liver parenchymal and non-parenchymal cell types. In the future, a combination of snRNAseq and scRNAseq may be needed to unlock the full liver cellular landscape and the impact of all liver cells for their ability to impact T cell-mediated alloreactivity in the liver.

In summary, scRNAseq analysis of pediatric liver allograft rejection reveals highly restricted CD8_EXP_ that retain an effector gene expression program and are detected within lobular areas of inflammation long after histologic resolution of rejection. Differential crosstalk between KC-CD8_EXP_ subpopulations reveals potential molecular targets which may drive tissue retention of CD8_EXP_ T cells, leaving patients at risk for rejection and allograft fibrosis when immunosuppression is weaned. These fundamental insights recapitulate recent findings in adult lung and kidney allograft rejection, paving the way for a cross-organ approach to develop targeted therapies for solid organ transplant rejection.

## Methods

### Sex as a biological variable

Male and female liver transplant recipients were included in the study, as detailed in Table 1. Due to the limited sample size as well as the complexity of receiving sex-mismatched allografts, we are unable to specifically determine the influence of sex on our results.

### Study Design

After obtaining informed consent/assent, children who received an isolated liver transplant at <17 years of age and who were currently being followed by the Liver Transplant Program at Cincinnati Children’s Hospital Medical Center, Cincinnati, OH were included in the study. Donor liver was obtained from a wedge liver biopsy obtained during backbench preparation of a donor liver prior to transplantation. No ACR is defined as not meeting any histologic criteria for ACR, which comprises a composite Rejection Activity Index (RAI or Banff score) of up to 9: portal inflammation (0–3), bile ductulitis (0–3), and venulitis (0–3). Late ACR was defined as occurring a minimum of 6 months after transplantation. For each study subject, the following data points were obtained from the electronic medical record: recipient and donor age/race/ethnicity/sex, indication for transplant, history of rejection prior to study entry, ALT, GGT, and tacrolimus trough level at the time of each biopsy, baseline immunosuppression, and medications used for treatment of rejection. Subjects were followed for a minimum of 2 years after rejection.

### Sample Collection

The diagnosis of rejection necessitates a liver biopsy. 16G core needle biopsies up to 2cm in length were obtained during a clinically-indicated non-targeted liver biopsy procedure to evaluate for ACR. Two biopsies were obtained from each subject. One biopsy was placed into 10% neutral buffered formalin and paraffin embedded (FFPE) according to standard procedure for clinical use. All treatment decisions were made independently by the practicing clinicians and not by the research study team. The FFPE biopsy samples corresponding to those which underwent scRNAseq were independently analyzed by a blinded pathologist for features of ACR (S. Ranganathan). The second biopsy sample, which was designated for research, was placed immediately into HypoThermosol FRS Preservative Solution (HTS) (BioLife Solutions Inc., cat #101102) at 4C for 15 minutes, then transferred to a cryovial containing cold CryoStor CS10 (BioLife Solutions, Inc., cat #210102) for 30 minutes on ice prior to freezing in a Mr. Frosty Freezing Container at −80C. Biopsy samples were transferred to liquid nitrogen within 24 hours and stored up to 3 years until analysis.

### Tissue Dissociation

The tissue dissociation protocol was modified from a previously described cold-active protease digestion originally developed for kidney biopsies (37, 71). Liver core biopsies were thawed, minced into 1-2mm pieces with a sharp scissors, and gently digested at 4C in RPMI+10% FBS containing 100mg/mL soy trypsin inhibitor (Roche, cat #10109886001), 250U DNase I (Roche, cat #4536282001), 5mM CaCl2, 25ug/mL collagenase A from clostridium histolyticum (Roche, cat #10103586001), and 25ug/mL collagenase type IV from clostridium histolyticum (Worthington, LS004186). Digestion was performed for 3min on ice with intermittent inversion and gentle pipetting with wide-bore pipet tips. The cell suspension was filtered via a pre-primed 30mm cell strainer, then centrifuged at 300g for 5 min at 4C. Trypan blue exclusion was used to determine cell viability and concentration. Cells were resuspended at 1,000cell/mL per the 10x Genomics Chromium protocol and immediately prepared for single-cell barcoding.

### Single-Cell barcoding, cDNA synthesis, and library preparation

Samples were processed for single-cell sequencing following the Chromium Next GEM Single Cell V(D)J Reagent Kit, version 1.1 protocol. Briefly, cells were uniquely barcoded by using 10x fluidics (10x Genomics Single Cell Controller) to combine each individual cell with an individual barcoded Single Cell 5’ Gel Bead creating a Gel Beads-in-Emulsion (GEMs) solution (10x Genomics, PN-1000165 and PN-1000020). An average of 17,400 cells were loaded to achieve an estimated recovery of 10,000 cells per biopsy. GEM gel beads were dissolved and cDNA was synthesized from the resulting tagged mRNA transcripts over 14 amplification cycles; 50ng of cDNA was used for the construction of each library. Total gene expression libraries (PN-10000020), enriched TCR libraries (PN-10000005) and enriched BCR libraries (PN-1000016) were created using the Single Index Kit T Set A (PN-10000213).

### Sequencing, alignment, and dataset processing

Total gene expression as well as TCR- or BCR-enriched libraries were sequenced on the NovaSeq 6000 sequencer using S2 flow cells, with the goal of obtaining >320M reads per sample. Raw sequencing data were processed using CellRanger software (version 6.0.0) following standard practices to generate FASTQ files (72). For the gene expression data, the ‘counts’ function was used with default parameters to generate a gene expression matrix. Similarly, the ‘vdj’ function was used with default parameters to call clonotypes for B cell and T cell receptors in the immunophenotyping data. The gene expression data for each dataset was filtered to retain cells with 250 or more features, 500 or more UMIs, and less than a 30% mitochondrial fraction. This resulted in a total of 10,566 cells for the 30 samples across 14 patients. Gene expression data for each sample was normalized using SCTransform (version 2) following the author’s instructions (73). After normalization, integration was performed using Seurat v4.0.3 as described by the authors using 3000 integration features, and clustering was done at the default resolution of 0.5, which resulted in 16 original clusters (74).

### Cell Classification

Dimensionality reduction plots were generated for each sample individually and for the integrated sample object to facilitate cell type classification. Differential expression to define cluster identities via Seurat was done on the ‘SCT’ assay slot as recommended by the authors using default parameters for minimum fraction expression and log fold change threshold parameters. Additionally, exploration of literature-derived marker genes and gene signatures for known liver and immune cell types were used to aid in cell type identification. Gene signature scoring was performed using custom scripts to define a score for expression of a set of genes in each cell relative to stably expressed control genes within the sample. Sub-clustering was done to resolve clusters containing mixed cell types that needed to be separated, including the T cell, endothelial cell, Kupffer cell, and B-cell compartments, which resulted in 22 distinct clusters that best reflected known cell types. These were further grouped into 12 broad cell type categories for visualization purposes. Barplots were created to convey proportion differences in the cell types with regard to histologic diagnosis, which was also overlaid onto TSNE plots. Similar plots were generated for T cell, B-cell, NK-cell, and Kupffer cell subsets. Dot plots were generated with Seurat using log-normalized expression values for visualization (scale = F).

### T Cell Receptor Analysis

Processing of the T cell (TCR) data from the Cell Ranger pipeline was done both individually for sample-level information and as an integrated analysis to incorporate into the existing GEX Seurat object. Metadata for clone size, TRA sequence, and TRB sequence from the individual analyses were added to the integrated object. UMAP and TSNE plots were generated with FeaturePlot in Seurat using “clonesize” as the feature for visualization. Additionally, UMAP and TSNE plots were created for each individual sample with “clonesize” as the visualized feature. TRA and TRB sequences were concatenated to assign a clonotype to each cell, which was then used to generate barcode frequency plots for each sample. Heatmaps were generated with the ‘pheatmap’ package for exploring shared clonotypes across samples and patients (75). Expanded clonotypes in this study were defined as those clonotypes present in three or more T cells within the same sample. Additionally, shared expanded clonotypes across biopsies within the same patient were visualized on connected barplots using the ‘immunearch’ package in R (76). Expression of marker genes for T cell activation, exhaustion, and residency were visualized with violin plots using Seurat.

### B Cell Receptor Analysis

Similar to the TCR analysis, processing of the B cell (BCR) data from the Cell Ranger pipeline was done both individually for sample-level information and as an integrated analysis to incorporate into the existing GEX Seurat object. Further, additional analysis of the immunophenotyping data for B cells was performed as described in Gill et al. (77), which broadly validated the Cell Ranger values for clone size and added hypermutation scores for each cell. These recalculated values were used for all downstream analyses. UMAP and TSNE plots were generated with FeaturePlot in Seurat to visualize clone size and hypermutation status. Finally, barplots were generated to show light chain and heavy chain usage per sample.

### Cell-Cell Interaction Prediction

Predictions for receptor-ligand interactions between clusters were performed using CellPhoneDB (78) with default parameters. Visualization of these interactions via chord diagrams was done using the ‘circlize’ and ‘chorddiag’ packages in R (79, 80). Expression of receptors and ligands from selected interactions, chosen from those interactions that differed between expanded and not-expanded T cell subsets, were visualized on TSNE plots. The table of shared interactions across cell types was generated in Excel using the relevant interactions tables.

### BaseScope Assay

Two sets of dual-index probes designed to bind to the hypervariable CDR3-a and CDR3-β regions of the most abundant T cell clone in Patient 6 and 13. 5um FFPE liver biopsy slides underwent deparaffinization, rehydration, treatment with protease IV, and antigen retrieval via boiling for 10 minutes according to the manufacturer’s instructions. Probes were hybridized to our target mRNA and amplified per the manufacturer’s instructions, with incubation extended at steps AMP7 and AMP11 from 30 to 60 minutes to improve probe detection. Slides were counterstained with 25% hematoxylin solution, washed with tap water, immersed in 0.02% ammonia water, and again washed with tap water before mounting. Brightfield images were captured at 40× magnification.

### Statistics

P-values less than or equal to 0.05 were considered statistically significant for all analyses. Differential cell type proportions were calculated using a Wilcoxon rank sum test. Student’s t-test was used to calculate differences between CD8_EXP_ T cell clonotype frequency and rejection subtype.

### Study Approval

This study was approved by the Cincinnati Children’s Hospital Institutional Review Board (IRB 2018-4660). Participants who were >18 years old at the time of enrollment provided written, informed consent. If participants were <18 years old at the time of enrollment, written informed consent was provided by one parent/guardian. Any study participant who turned 18 during the 2 year study follow up period was re-consented at that time. Informed consent was provided before any study procedures occurred.

## Data Availability

All data used in this paper have been deposited in the Gene Expression Omnibus (GEO) with the GEO accession number to be made publicly available upon manuscript acceptance. The scripts used to generate the data and figures presented in this paper can be found at the project’s GitHub repository: https://github.com/EDePasquale/Peters-single-cell.

## Author Contributions

ALP, EAKD, DAH, and ESW designed the research studies. ALP and GB conducted experiments. ALP recruited participants. EAKD and KMR analyzed data. EAKD and ALP wrote the manuscript. ALP, EAKD, GB, KMR, ESW, and DAH edited the manuscript. ALP and EAKD are co-first authors for this manuscript, however ALP and her research lab initiated the project and is therefore listed first.

## Acknowledgements

We would like to thank all of the study participants and their families for contributing to make this research possible. This project was supported, in part, by NIH P30 DK078392 Integrated Research Pathology Core (Dr. Ranganathan, SCR_022637), Single Cell Genomics Facility (Kelly Rangel, Shawn Smith, RRID: SCR_022655), Research Flow Cytometry Core (Dr. Celine Silves-Lages, SCR_022628), and Bio-Imaging and Analysis Facility (Dr. Matthew Kofron, SCR_022628), of the Digestive Diseases Research Center in Cincinnati, OH and Cincinnati Children’s Hospital Research Foundation. A.L.P. was supported by NIH 5K12HD028827-28/29, the American Society of Transplantation Career Transition Research Award, a Digestive Health Center Pilot and Feasibility Award (CCHMC; NIH P30 DK078392), the Markham Family Liver Transplant Research Award (CCHMC), and donations from the Weidner family.

---

**Supplemental Figure 1.**
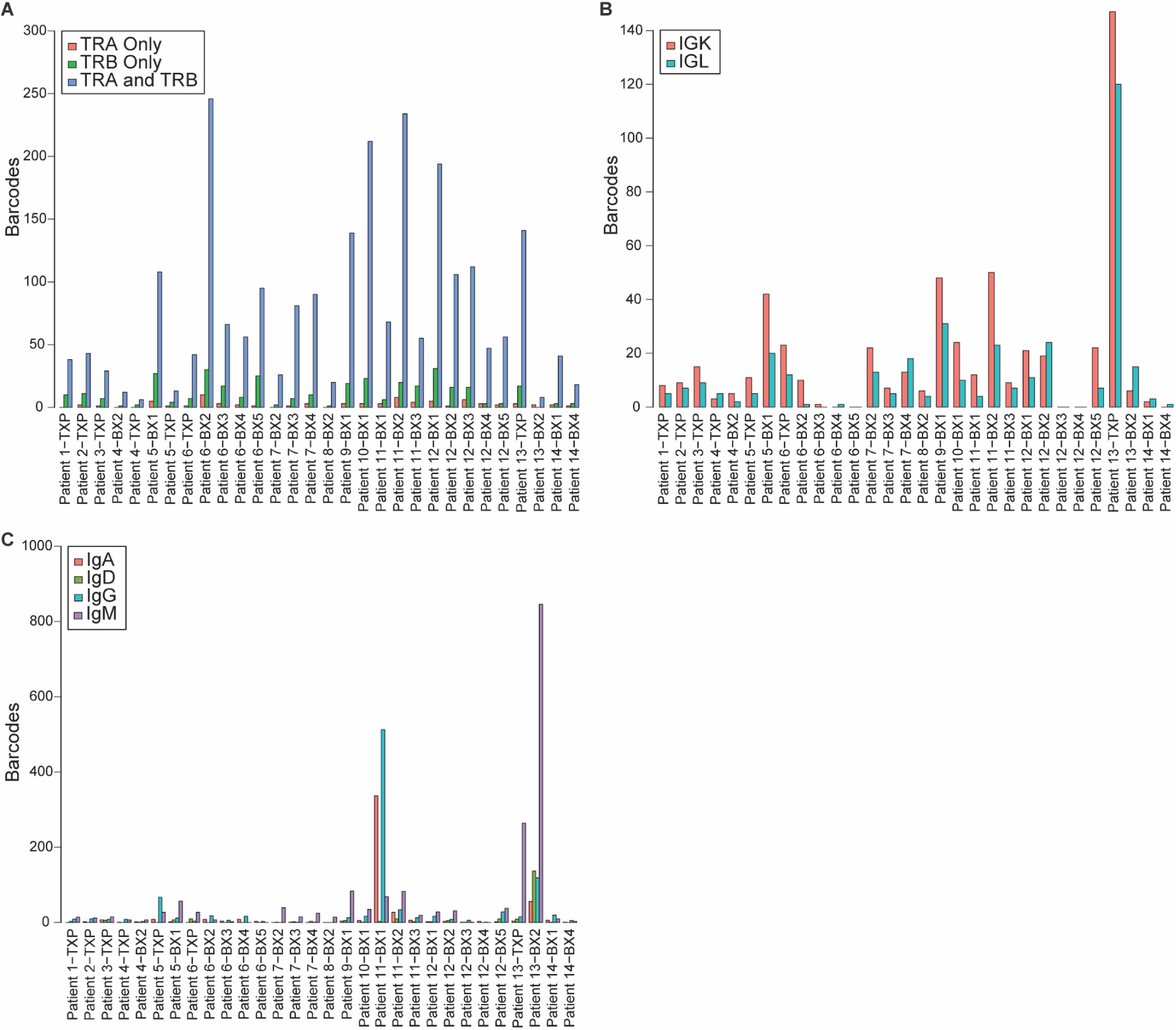
TRA/B and Immunoglobulin locus usage by sample. Grouped bar plot showing for each sample (X-axis) the number of cells containing: (**A**) TRA only (red), TRB only (green), or both TRA and TRB (blue) genes (Y-axis), (**B**) IGK (Immunoglobulin kappa light chain; red) or IGL (immunoglobulin lambda light chain; teal)). (**C**) Distribution of cells expressing heavy chain genes IgA, IgD, IgG, or IgM.

**Supplemental Figure 2.**
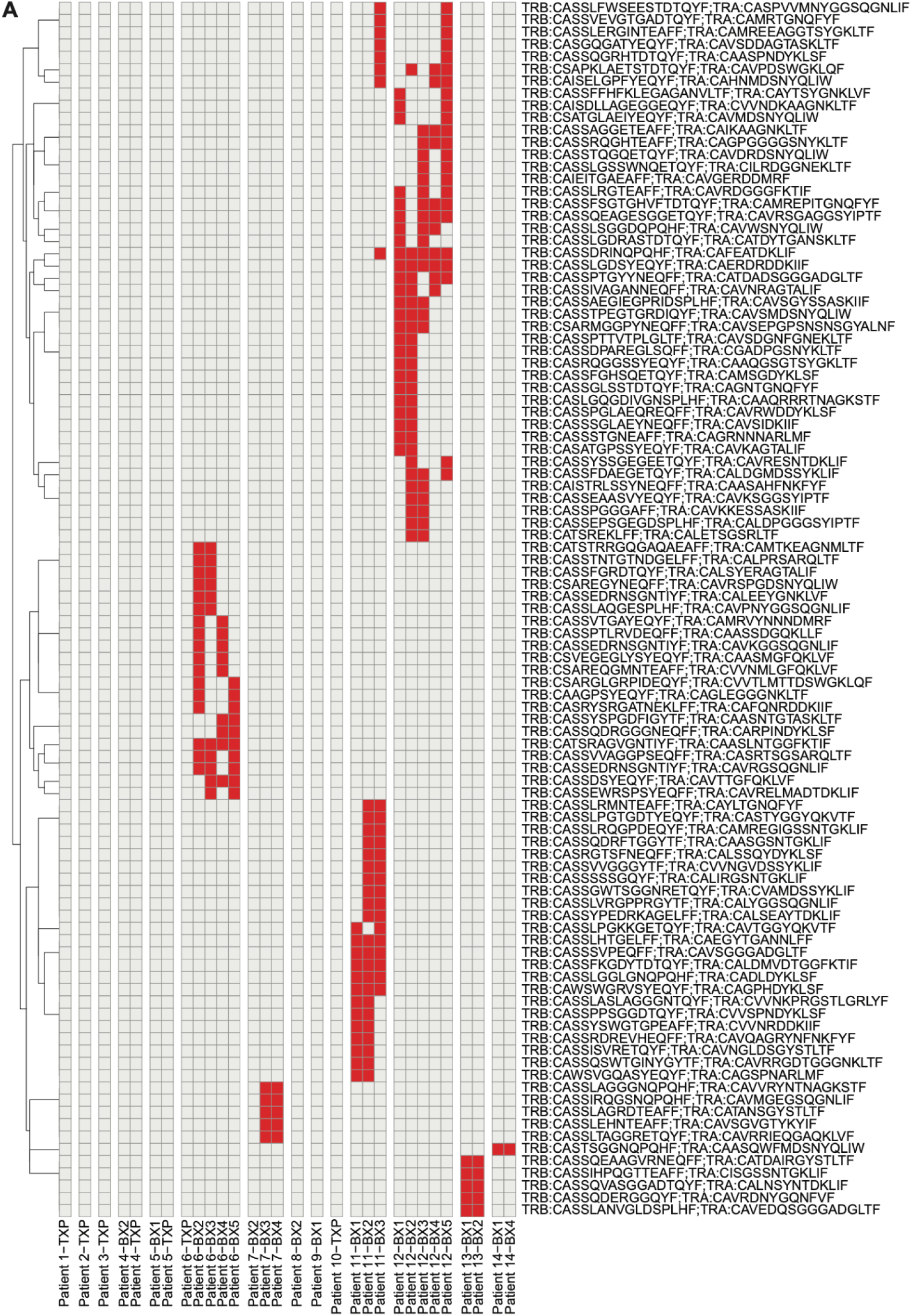
Shared expanded clonotypes across samples. (**A**) Heatmap showing clonotypes (rows) that are shared across samples (columns), with columns grouped by patient. Red indicates a shared expanded clonotype while gray indicates that the clonotype was not shared between samples.

**Supplemental Figure 3.**
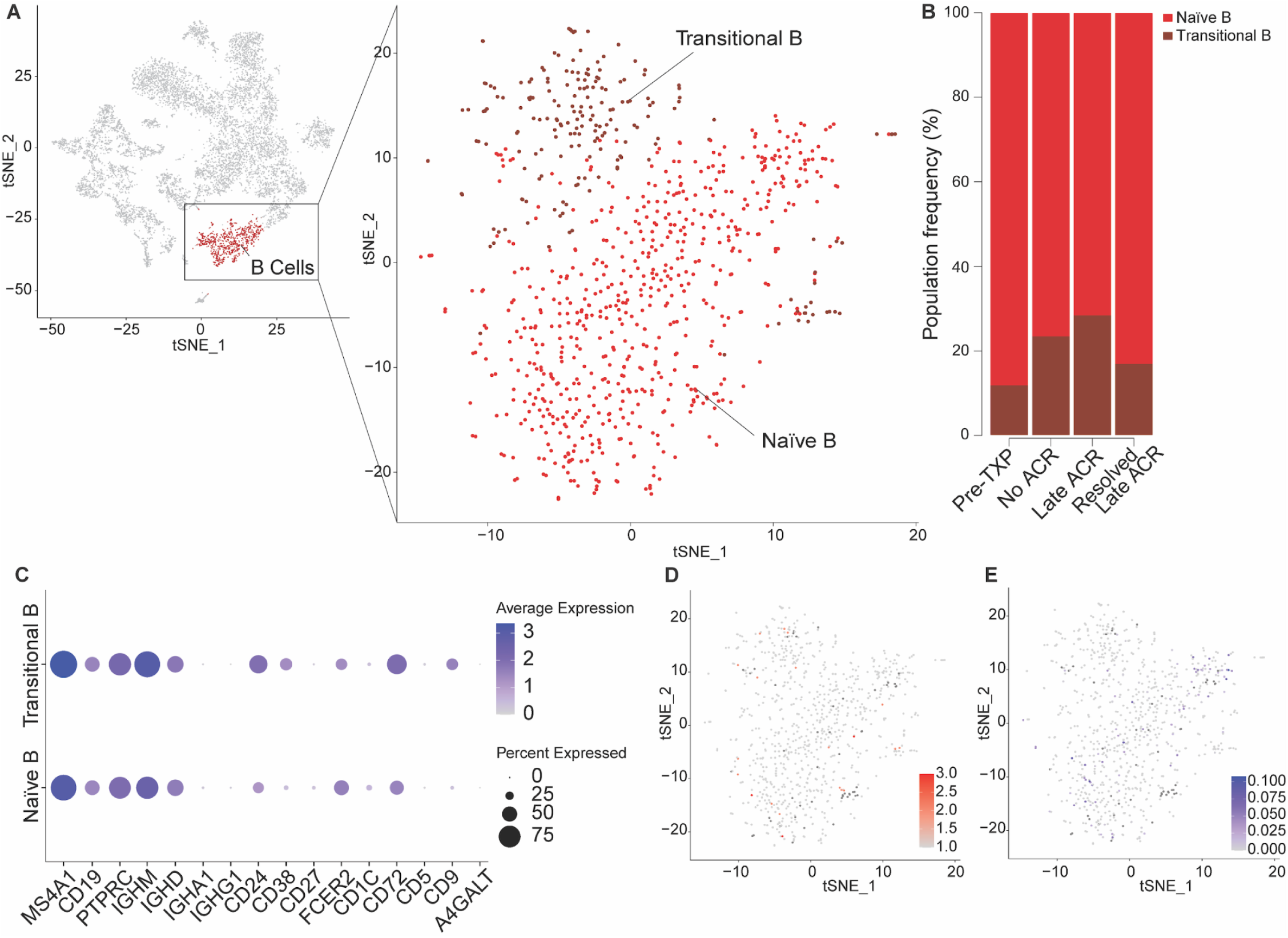
B-cell subtype identification. (**A**) B-cells were isolated and clustered separately to define additional subpopulations of cells and visualized on a TSNE plot. (**B**) Stacked bar plots indicating B-cell subtype frequency for each histologic diagnosis. (**C**) Dot plot showing average expression for key B-cell defining genes (indicated by color) and the percent of cells within each B-cell subtype that express that gene (indicated by dot size). (**D**) TSNE plot of B-cells colored by clone size, with light gray indicating no clonal expansion, bright red indicating high clonal expansion, and dark gray indicating no B-cell receptors detected. (**E**) Estimates of somatic hypermutation levels for each cell, with dark blue indicating more hypermutation and light gray indicating less. Abbreviations: Acute cellular rejection (ACR).

**Supplemental Figure 4.**
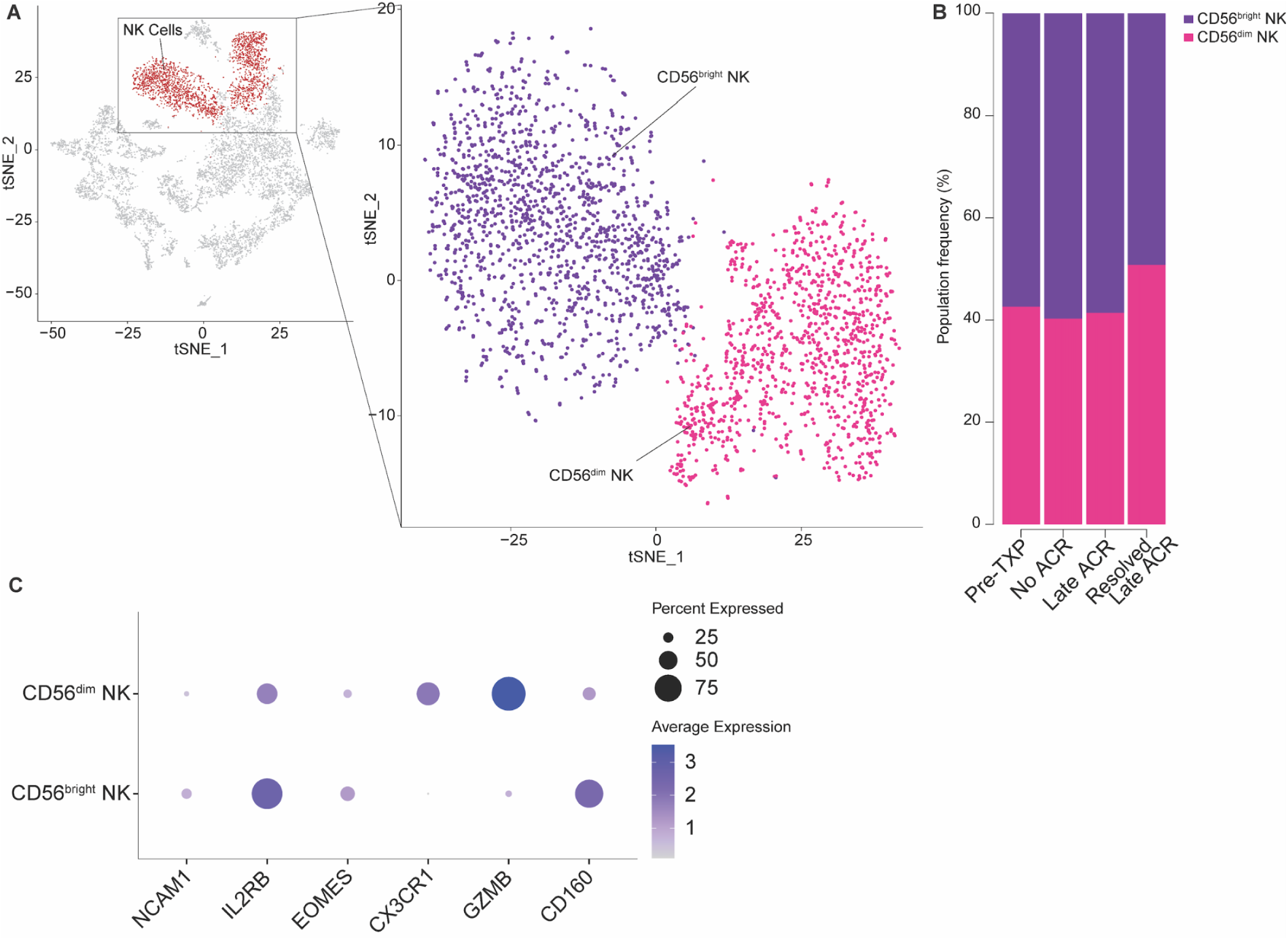
NK-cell subtype identification. (**A**) Natural killer cells were isolated and sub-clustered to define two clear subtypes, CD56^bright^ NK and CD56^dim^ NK, and visualized on a TSNE plot. (**B**) Stacked bar plot to indicate the proportion of each NK sub-cluster within all NK cells. (**C**) Dot plot showing average expression of key identifying genes for each sub-cluster. Abbreviations: Natural killer (NK).

**Supplemental Figure 5.**
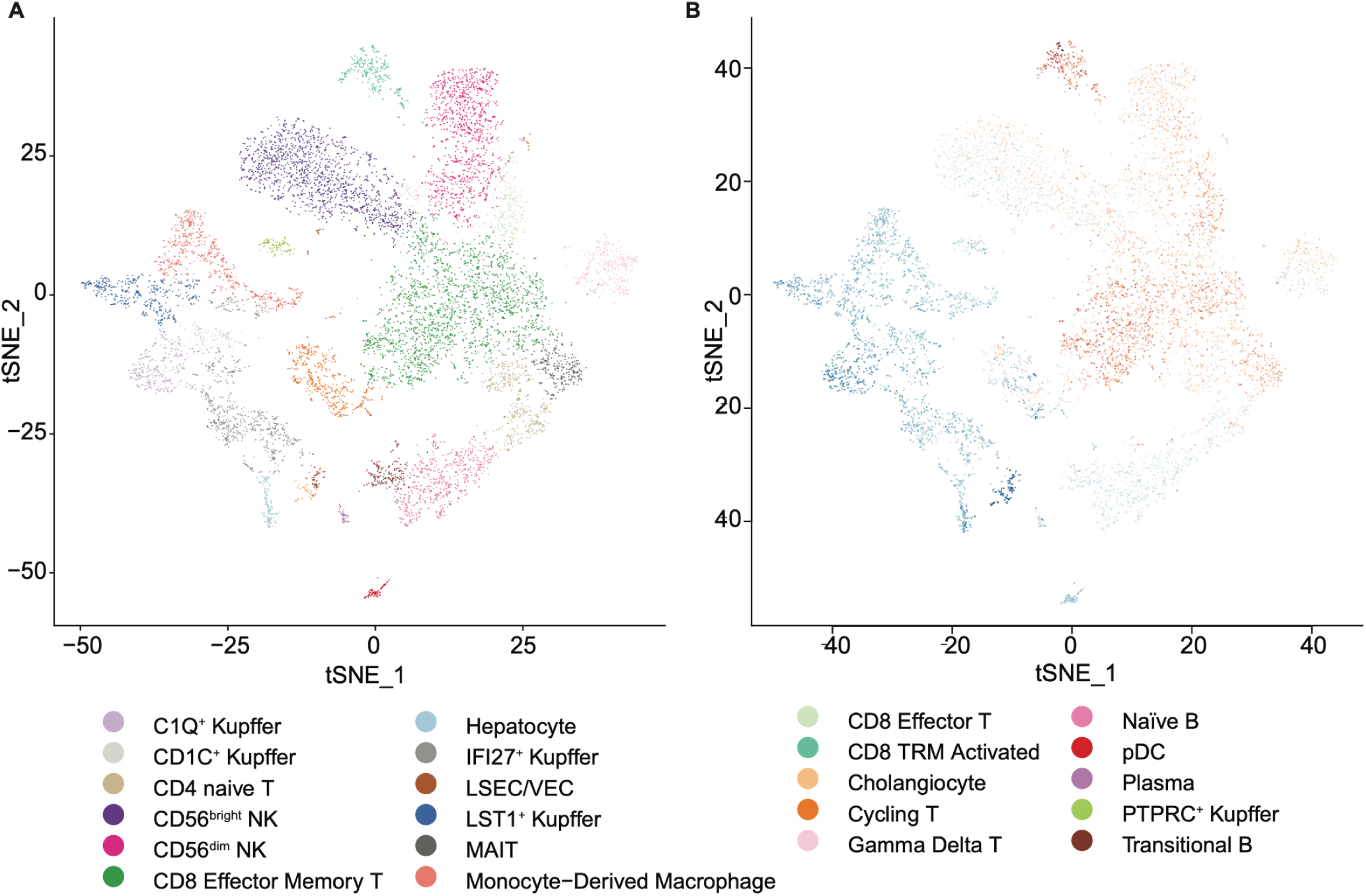
(**A**) TSNE plot colored by high resolution clusters for all cell types. (**B**) TSNE plot of composite expression of the 200-gene rejection-associated signature derived from the iWITH trial, showing increased expression in T and NK subpopulations within the scRNAseq dataset. Blue=low expression, Red = high expression.

## Notes

**Conflict of Interest**: The authors have declared that no conflict of interest exists.

### Competing Interest Statement

The authors have declared no competing interest.

